# Spatial modeling reveals nuclear phosphorylation and subcellular shuttling of YAP upon drug-induced liver injury

**DOI:** 10.1101/2022.03.31.486549

**Authors:** Lilija Wehling, Liam Keegan, Paula Fernández-Palanca, Reham Hassan, Ahmed Ghallab, Jennifer Schmitt, Peter Schirmacher, Ursula Kummer, Jan G. Hengstler, Sven Sahle, Kai Breuhahn

**Affiliations:** Institute of Pathology, University Hospital Heidelberg, Germany; Department of Modeling of Biological Processes, COS Heidelberg/BioQuant, Heidelberg University, Germany; Leibniz Research Centre for Working Environment and Human Factors, Department of Toxicology, Technical University Dortmund, Germany; Institute of Biomedicine (IBIOMED), University of León, Campus de Vegazana s/n, 24071 León, Spain; Centro de Investigación Biomédica en Red de Enfermedades Hepáticas y Digestivas (CIBERehd), Instituto de Salud Carlos III, Av. de Monforte de Lemos 5, 28029 Madrid, Spain; Department of Forensic Medicine and Toxicology, Faculty of Veterinary Medicine, South Valley University, 83523, Qena, Egypt

**Keywords:** DILI, computational modeling, partial differential equations (PDE), Hippo pathway, acetaminophen (APAP)

## Abstract

The Hippo signaling pathway controls cell proliferation and tissue regeneration *via* its transcriptional effectors *yes-associated protein* (YAP) and *transcriptional coactivator with PDZ-binding motif* (TAZ). In this context, the canonical pathway topology is characterized by sequential phosphorylation of kinases in the cytoplasm that define the subcellular localization of YAP and TAZ. However, the molecular mechanisms controlling the nuclear/cytoplasmic shuttling dynamics of both factors under physiological and tissue-damaging conditions are poorly understood. By implementing experimental data, *partial differential equation* (PDE) modeling, as well as automated image analysis, we demonstrate that nuclear phosphorylation contributes to differences between YAP and TAZ localization in the nucleus and cytoplasm. Treatment of hepatocyte-derived cells with hepatotoxic *acetaminophen* (APAP) overdose induces a biphasic protein phosphorylation eventually leading to nuclear protein enrichment of YAP but not TAZ. APAP-dependent regulation of nuclear/cytoplasmic YAP shuttling is not an unspecific cellular response but relies on the sequential induction of *reactive oxygen species* (ROS), *RAC-alpha serine/threonine-protein kinase* (AKT, synonym: *protein kinase B*), as well as elevated nuclear interaction between YAP and AKT. Mouse experiments confirm this consecutive sequence of events illustrated by the expression of ROS-, AKT-, and YAP-specific gene signatures upon APAP administration. In summary, our data illustrate the importance of nuclear processes in the regulation of Hippo pathway activity. YAP and TAZ exhibit different shuttling dynamics, which explains distinct cellular responses of both factors under physiological and tissue-damaging conditions.

**Significance:** We show that canonical view on the Hippo pathway must be extended by additional regulatory processes in cell nuclei. These processes significantly contribute to the activity of YAP and TAZ under unchallenged conditions (e.g., with cell density as physiological regulator of the Hippo kinase cassette) or under cell damaging conditions (e.g., after administration of APAP overdose). APAP-induced cellular damage activates YAP *via* distinct molecular processes as part of a cell-protective response.

## Introduction

The evolutionary conserved Hippo signaling pathway controls tissue homeostasis through the regulation of cell proliferation, apoptosis, differentiation, and cellular fate (Zhao et al., 2011). This pathway is regulated by extracellular information derived from extracellular matrix stiffness, actomyosin dynamics and cell density (Deng et al., 2015; Zhao et al., 2007), which in turn modulate a serine/threonine kinase cassette consisting of *sterile 20-like kinase 1/2* (MST1/2), and *large tumor suppressor 1/2* (LATS1/2). According to the canonical Hippo pathway model, active cytoplasmic LATS1/2 phosphorylate and inactivate two important downstream pathway effectors: the transcriptional co-activators *yes-associated protein* (YAP) and its paralog *transcriptional coactivator with PDZ-binding motif* (TAZ). This phosphorylation is associated with nuclear YAP/TAZ exclusion followed by their proteasomal degradation. In contrast, Hippo pathway inactivation in the cytoplasmic compartment causes YAP/TAZ dephosphorylation and nuclear translocation. In the nucleus, YAP/TAZ bind DNA sequence-specific transcription factors such as *TEA DNA-binding proteins* (TEADs) or *forkhead box protein M1* (FOXM1), which control the transcription of genes involved in e.g., cell cycle control and paracrine communication (Marti et al., 2015; Thomann et al., 2020; Weiler et al., 2017).

Due to its pivotal role in the regulation of cell proliferation, the Hippo/YAP/TAZ axis is important for tissues maintenance during regenerative processes. Indeed, YAP- or YAP/TAZ-deficiency reduce the regenerative capacity of tissues such as liver, skin, and heart (M.-J. Lee et al., 2014; Lu et al., 2018; Xin et al., 2013). As exemplified in detail for the liver, YAP is activated in hepatocytes under disease conditions that support a continuous regenerative response in different *in vivo* model systems (Machado et al., 2015; Mooring et al., 2020). Interestingly, the roles of YAP and TAZ are not identical, since TAZ, but not YAP, contributes to fat accumulation in the liver (steatosis) (X. Wang et al., 2016). In contrast, YAP protects from liver ischemia-reperfusion injury by promoting tissue repair (Liu et al., 2019). These findings strongly suggest a cell-protective role of the Hippo pathway under liver injury conditions; however, the role of YAP and TAZ differ. As YAP and TAZ control the expression of similar target genes in different cell types (Weiler et al., 2020; Zanconato et al., 2015), other regulatory mechanisms must account for these phenotypic differences such as differential nuclear/cytoplasmic shuttling (Reggiani et al., 2021). However, detailed comparative studies regarding subcellular localization dynamics for YAP and TAZ are missing as well as spatially resolved computational framework such as *partial differential equation* (PDE) models.

Recent data demonstrated a direct impact on YAP by *acetaminophen* (APAP), which is a widely used analgesic drug and leading cause of acute liver failure (Poudel et al., 2021). Here, APAP overdose in mice caused liver injury, which was associated with a prominent nuclear YAP enrichment in hepatocytes. Interestingly, silencing of YAP did not impair liver regeneration (which would be expected after inactivation of this pro-proliferative factor), but promoted tissue regeneration upon APAP-induced liver damage (Poudel et al., 2021). In contrast, a supportive role of YAP in tissue repair and regeneration was suggested by chemical blockade of upstream Hippo pathway kinases MST1/2, which reverted APAP-induced liver damage (Fan et al., 2016). These results point to intricate and context-specific roles of YAP and/or TAZ under regenerative conditions.

Our study comprises a quantitative and mechanistic analysis of YAP/TAZ dynamics based on spatially resolved, high-throughput confocal live-cell imaging data and PDE modeling. By using this experimental and computational toolbox, we show that nuclear phosphorylation of YAP/TAZ is important for their subcellular shuttling dynamics and that both factors do not equally respond to cell density and APAP-induced hepatocellular damage. We mechanistically show that APAP overdose stimulates a rapid *reactive oxygen species* (ROS) response, which facilitates the physical nuclear interaction of YAP and AKT as predicted by computational modeling. Thus, our data demonstrates for the first time that APAP-induced YAP activation is a dynamic and specific process that relies on the sequential activation of nuclear molecular events.

## Results

### Establishment of an in vitro model for measuring time-resolved spatial localization of YAP and TAZ in hepatocellular cells

Due to technical limitations, primary human hepatocytes are not suitable for the analysis of dynamic YAP/TAZ shutting *in vitro*: they rapidly undergo trans-differentiation in culture and different human donors exhibit genomic/epigenetic variability, which diminish reproducibility. We therefore selected the hepatocyte-derived liver cancer cell line Hep3B that was characterized by a prominent YAP/TAZ and cytochrome P450 2E1 (CYP2E1) expression (Supplementary Figure S1A) (Weiler et al., 2020). Detectable CYP2E1 levels are required for the enzymatic transformation of APAP to toxic *N-acetyl-p-benzoquinone imine* (NAPQI) (S. S. Lee et al., 1996).

Since we were interested in cell-to-cell variability and spatial localization information of YAP and TAZ, we established a cell line that allowed the quantitative and spatio-temporal investigation of subcellular YAP/TAZ distribution near the single cell resolution. For this, we generated Hep3B cells that stably express Venus-tagged YAP (mVenus-YAP) and mCherry-tagged TAZ (mCherry-TAZ) (Figure 1A-B). In addition, mCerulean-tagged histone H2B (mCerulean-H2B) allowed spatial allocation of YAP/TAZ. These chosen fluorophores facilitated optimal discrimination between emission spectra by live cell microscopy (Supplementary Figure S1B). Subcellular localization changes of tagged YAP and TAZ under variable cell density conditions illustrated functionality of both proteins regarding nuclear-cytoplasmic shuttling (Figure 1B). However, we observed a high degree of intracellular variability.

**Figure 1.**
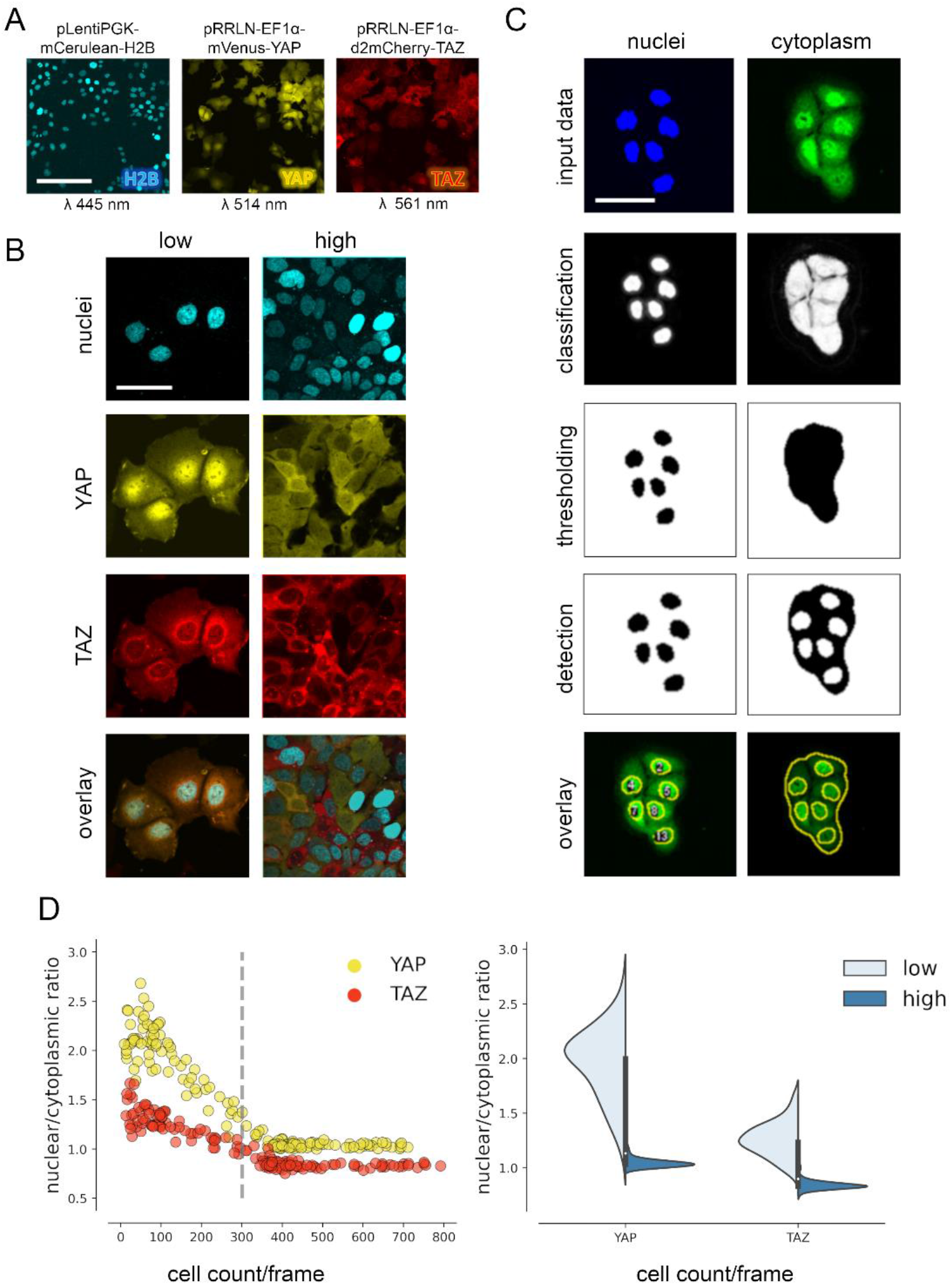
YAP and TAZ show distinct nuclear shuttling in hepatocellular cells upon increasing cell density. **A.** The hepatocyte-derived cell line Hep3B was transduced using three lentiviral vectors coding for mCerulean-tagged H2B (pLentiPGK-mCerulean-H2B), mVenus-tagged YAP (pRRLN-EF1α-mVenus-YAP) and mCherry-tagged TAZ (pRRLN-EF1α-d2mCherry-TAZ). Combined treatment with the antibiotics hygromycin, geneticin, and blasticidin selected for triple-positive cells. Cells were confocally imaged in three channels: 445 nm (CFP, Cerulean), 514 nm (YFP, Venus) and 561 nm (RFP, mCherry). Scale bar: 250 µm. **B.** Exemplary pictures illustrating the subcellular localization of H2B, YAP, and TAZ proteins under low (left) and high (right) cell density conditions. Scale bar: 50 µm. **C.** Automatic image analysis workflow depicts analysis of nuclear (left) and cytoplasmic (right) fluorescence intensity in confocal images of living cells. The acquired images were pre-screened for imaging artifacts (e.g., out-of-focus images were excluded) and subjected to the image analysis pipeline in Fiji. Object classification (Weka algorithm), thresholding and object counting were performed. Object masks were overlaid with original data and the NCR was calculated by dividing average fluorescent signal intensity of YAP/TAZ in the nucleus with the fluorescence in the cytoplasm. Scale bar: 50 µm. **D.** Quantification of YAP and TAZ NCR under increasing cell density conditions. One of four representative experiment is depicted. *Left*: the NCR of YAP (yellow, n=310) and TAZ (red, n=310) were plotted against cell count per visual field (0.4 mm^2^). A high NCR value indicate predominant nuclear localization. Dashed line shows mean cell count. *Right*: violin plots summarize shift in NCR between low (below mean cell count) and high (above mean cell count) cell density for YAP and TAZ.

For analysis of variable high-throughput data, we developed an algorithm for the semi-quantitative measurement of tagged YAP and TAZ in nuclei and cytoplasm of cells. In brief, spatial image data was classified based on the expression of Venus-YAP and mCherry-TAZ using the Weka segmentation algorithm (Figure 1C and Supplementary Figure S1C) (Arganda-Carreras et al., 2017). The fluorophore intensity measurements were used to calculate the *nuclear/cytoplasmic ratio* (NCR), which characterized the relative nuclear enrichment of YAP or TAZ. Therefore, this approach allowed us to simultaneously define slightest subcellular changes of YAP and TAZ in small cell populations (field of view) as well as individual cells under live-cell conditions.

Subsequently, the fluorophore-expressing cells were grown under variable cell density conditions and the YAP/TAZ localization was measured by confocal microscopy. The results confirmed that tagged YAP and TAZ dynamically shuttled between cytoplasm and the nucleus depending on amount of cell-cell contacts (cell density) (Figure 1D). However, TAZ showed a considerably less pronounced dynamic behavior compared to YAP (Figure 1D). YAP was predominantly detectable in cell nuclei under low cell density conditions (NCR>1), while increasing cell density led to cytoplasmic enrichment of both factors (NCR<1). Using this algorithm, cell density-dependent shuttling was equally demonstrated for other liver cancer cells (Tóth et al., 2021).

Together, the confocal imaging and image processing pipeline efficiently detects subtle differences in the dynamic subcellular changes of YAP and TAZ.

### Mathematical modeling predicts nuclear phosphorylation of YAP and TAZ

To investigate why Hippo pathway effectors YAP and TAZ differently respond to a low cell density environment and which Hippo pathway topology can explain the observed localization differences, we mathematically modelled the Hippo pathway using a partial differential equation (PDE) modeling framework (for detailed description of the PDE modeling approach refer to Materials and Methods and supplementary material).

First, we investigated a canonical Hippo pathway model, where phosphorylation takes place in the cytoplasm and where un-phosphorylated YAP/TAZ shuttle to the nucleus (Figure 2A). However, this mathematical model was not able to recapitulate the localization patterns which were observed in the experimental data (Figure 2B). For example, this model cannot sufficiently explain the homogenous nuclear dissemination of YAP or the distribution of TAZ at the nuclear membrane. Moreover, the residuals (experimental data minus model simulation) indicate that the largest discrepancies between model prediction and experimental data lie in the perinuclear cell area. Therefore, we considered alternative model topologies, which could explain the experimental data better.

**Figure 2.**
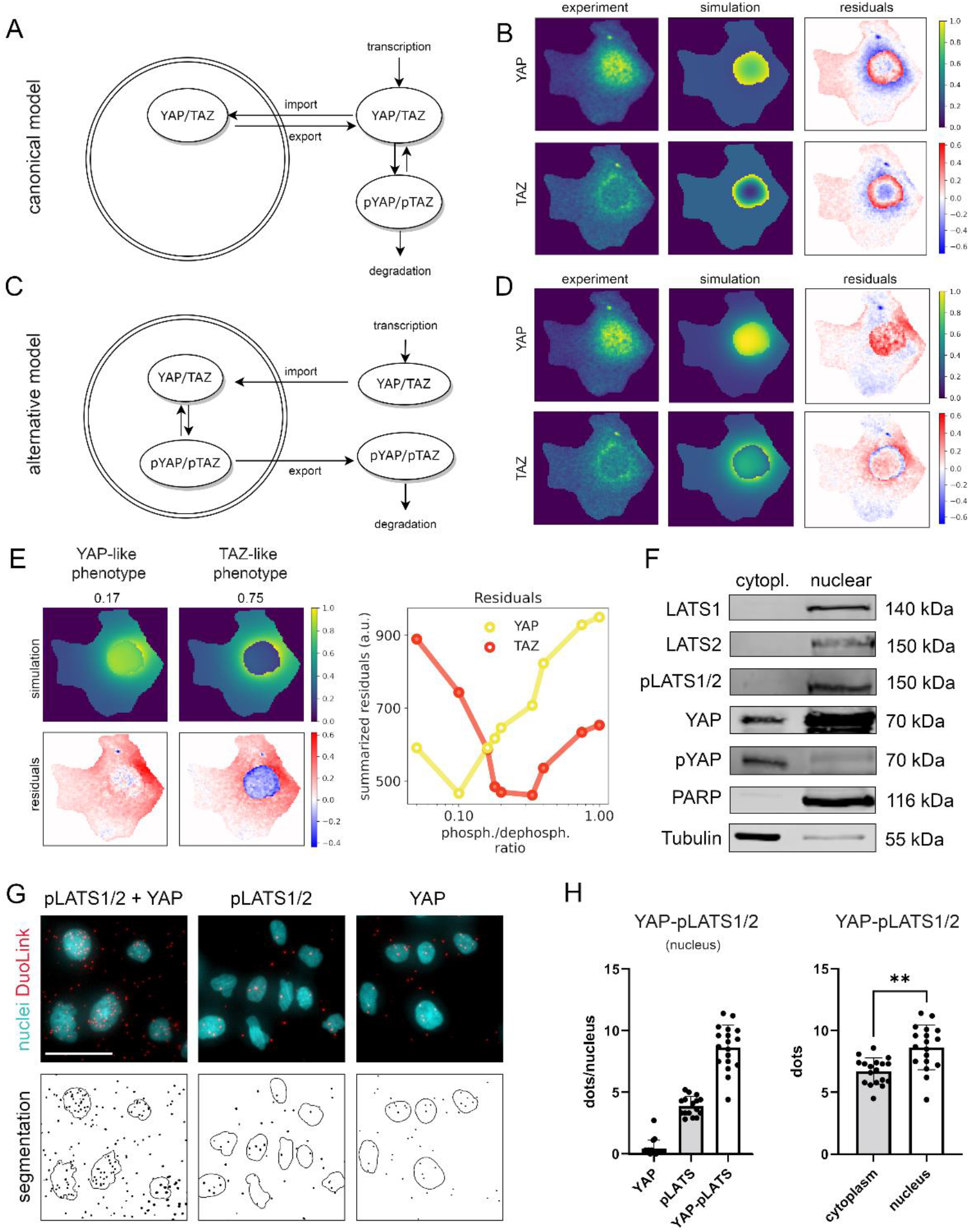
Mathematical modeling predicts that nuclear phosphorylation controls YAP/TAZ subcellular localization. **A.** The model reaction scheme of the canonical Hippo pathway. Only un-phosphorylated YAP/TAZ can enter the nucleus, whereas pYAP and pTAZ are exclusively localized to the cytosol. The phosphorylation of YAP/TAZ takes place exclusively in the cytosol. **B.** PDE model simulation of the canonical Hippo pathway compared to the experimentally measured YAP/TAZ localization. The residuals (experimental data minus model simulation) indicate low spatial accordance of the model simulation with the experimentally measured subcellular localization of YAP/TAZ. Images were normalized to the maximal value within each image. **C.** The model reaction scheme of the alternative Hippo pathway model. Phosphorylation and de-phosphorylation of YAP/TAZ takes place in the nucleus. Un-phosphorylated YAP/TAZ is imported in the nucleus and phosphorylated YAP/TAZ is rapidly exported to the cytoplasm. **D.** PDE simulation of the alternative model compared to experimental data. Residuals for the subcellular localization (e.g., nuclear distribution) sufficiently reflects results from confocal microscopy. **E.** Simulated impact of the nuclear phosphorylation to dephosphorylation ratio on subcellular localization of YAP in the alternative Hippo pathway model. *Left*: model simulation of two phosphorylation/dephosphorylation rates (0.17 and 0.75). *Right*: summarized residuals with respect to experimental data of YAP and TAZ as a function of phosphorylation to dephosphorylation ratio in the nucleus. **F.** Western immunoblot after nuclear and cytoplasmic protein fractionation under low cell density conditions. The central Hippo pathway kinases LATS1/2 are exclusively localized in the nuclear fraction. As expected, high pYAP levels are detectable in the cytoplasm, illustrating that the protein is transported outside the nucleus upon phosphorylation. PARP and Tubulin serve as loading controls for nuclear and cytoplasmic fractions, respectively. Equal amounts of cytoplasmic and nuclear proteins were loaded (n=4; one representative experiment shown). **G.** PLA for phosphorylated LATS1/2 (pLATS1/2) and YAP *(top row*). Red dots indicate physical interaction between pLATS1/2 and YAP. DAPI-stained nuclei are indicated in cyan. Individual pLATS and YAP antibodies serve as assay controls. *Bottom row*: computational segmentation of nuclei (empty circles) and dots (black spots). Scale bar: 50 µm. **H.** Quantification of the PLA assay. Each dot represents an individual image. *Left*: quantification of interactions between YAP and pLATS1/2 in the nucleus (n=18) compared to the negative controls (pLATS1/2 and YAP antibodies alone, n=16 and n=17, respectively). *Right*: The number of PLA dots of pLATS1/2 and YAP interaction in cytoplasm and in nuclei (n=18). Statistical test: two tailed paired t-test (p-val=0.005). For more information see Supplemental Figure S3 and Materials and Methods.

We generated an alternative PDE model, which precisely described YAP/TAZ localization as observed in the image data. In this model, a phosphorylation and de-phosphorylation reaction of YAP and TAZ in the nucleus was included (Figure 2C-D, Supplementary Figure S2A-B). Moreover, the alternative model describes that un-phosphorylated YAP/TAZ are transported to the nucleus and pYAP/pTAZ are rapidly excluded from the nucleus, which agrees with the general consensus that phosphorylated YAP and TAZ predominantly localize in the cytoplasm (Supplementary Figure S2C).

Since YAP and TAZ are considered biochemically similar molecules the question arises how the pronounced observed differences in the spatial distribution patterns can be explained. Using the alternative Hippo pathway model we demonstrated that the different phenotypes could be reproduced with one single PDE model where only the parameters of the phosphorylation and dephosphorylation reactions for YAP/TAZ in the nucleus differed, while all other parameters (including nuclear import and export reactions) were kept the same (Figures 2E, Supplementary Figure S2D).

In detail, low nuclear phosphorylation/dephosphorylation ratio (0.17) induced nuclear localization of YAP, while high ratio (0.75) induced nuclear protein exclusion. Upon increased phosphorylation activity (>0.2), the nuclear fraction of TAZ decreased and the summarized residual term for TAZ was reduced. According to residual terms, best accordance between the model and the experimental data was achieved at low phosphorylation/dephosphorylation ratio for YAP and at higher phosphorylation/dephosphorylation ratio for TAZ (Figure 2E, right panel). These findings suggest that differences in the nuclear phosphorylation rates of YAP and TAZ can explain the distinct shuttling patterns of both factors (as observed in Figure 1B). In addition, TAZ may require stronger phosphorylation than YAP to induce its efficient shuttling.

The results of the alternative mathematical model required that major components for YAP/TAZ phosphorylation process were present in the cell nucleus. Indeed, subcellular fractionation experiments illustrated that the Hippo pathway kinases LATS1/2 (total and phosphorylated) were predominantly localized in nuclear protein fraction (Figure 2F). In addition, a *proximity ligation assay* (PLA) illustrated that the pLATS1/2 physically interacted with YAP not only in the cytoplasm but also in the nucleus (Figure 2G). To quantitatively compare the number of YAP/pLATS interactions in cell nuclei and cytoplasm, we utilized a computational algorithm counting signals in nuclear and cytoplasmic area of the cells (Supplementary Figure S3). The signal quantification confirmed that YAP and pLATS interactions were detectable in both compartments with significantly higher interaction count/intensity in cell nuclei (Figure 2H).

In summary, our findings suggest that nuclear YAP/TAZ phosphorylation contributes to their inactivation and nuclear exclusion of YAP and TAZ. Moreover, variable phosphorylation rates of YAP and TAZ explain the subcellular shuttling differences between both factors.

### APAP regulates YAP protein localization and activity

To test if our findings are of relevance under conditions where YAP and TAZ are actively regulated, we decided to use a *drug-induced liver injury* (DILI) model. For this, we treated Hep3B cells expressing tagged mVenus-YAP and mCherry-TAZ with APAP (10 nM) (Barbier-Torres et al., 2017), and quantitatively investigated the dynamic shuttling of both factors.

Image quantification revealed that APAP led to a gradual and time-dependent nuclear enrichment of YAP after 48 hours, while TAZ only weakly responded to APAP (Figure 3A-B). This APAP-induced nuclear shuttling effect was clearly detectable under high cell density culture conditions, which is characterized by nuclear YAP/TAZ exclusion in untreated cells. No obvious response was detectable for YAP and TAZ at earlier time points post APAP treatment (Supplementary Figure S4A-B).

**Figure 3.**
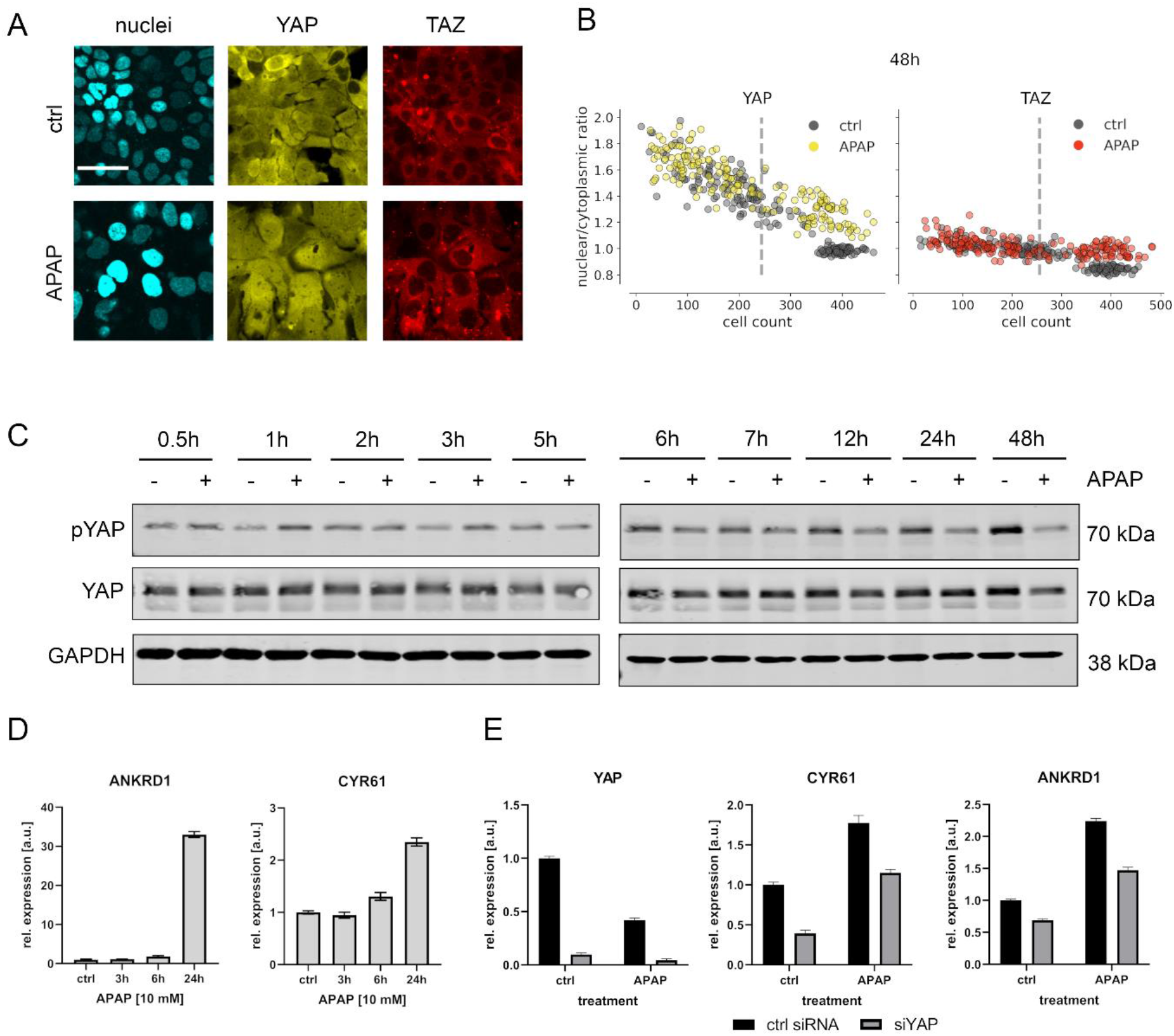
APAP regulates YAP phosphorylation, localization, and expression of its target genes. **A.** Live cell confocal imaging of H2B-cerulean, YAP-Venus and TAZ-mCherry in Hep3B cells. *Upper row:* control treatment (PBS). *Lower row*: APAP treatment (10 mM) induces nuclear enrichment of YAP, but not TAZ protein after APAP treatment within 48 hours (n=4; one representative experiment shown). Scale bar: 50 µm. **B.** Live cell confocal microscopy image quantification. NCR of YAP and TAZ with APAP (10 mM for 48 hours; yellow and red circles, n=180 per channel) and control (PBS; black circles, n=180 per channel) treatment under increasing cell density conditions (represented as mean cell count per visual field). Each dot represents a single visual field. Dashed line: mean cell density. **C.** Western immunoblot of the time-dependent protein abundancy of YAP and pYAP after APAP treatment (10 mM) in Hep3B cells (n=6; one representative experiment shown). GAPDH served as a loading control. **D.** Relative expression of the YAP target genes ANKRD1 and CYR61 24 hours after APAP (10 mM) treatment in HLF cells (n=2; one out of two biological replicates shown). **E.** Rescue experiment: relative expression of YAP and its target genes CYR61 and ANKRD1 with and without siRNA-mediated YAP inhibition (for 24 hours) with or without APAP (10 mM) treatment for 24 hours (n=2; one out of two biological replicates shown).

The APAP-induced nuclear enrichment of YAP was confirmed with cell population-based Western immunoblotting followed by detection of total and phosphorylated YAP (Figures 3C, Supplementary Figure S4C). Notably, at early time points YAP hyperphosphorylation indicated protein inactivation (up to 3 hours after APAP administration). However, between 24 to 48 hours after APAP treatment, a clear YAP de-phosphorylation was detectable (Figure 3C). Similar results were observed for another hepatocyte-derived cell line HepG2 (Supplementary Figure S4C).

To test if APAP-dependent de-phosphorylation/activation of YAP at later time points is leading to its transcriptional activation, we measured the relative expression of known YAP target genes *cysteine rich angiogenic inducer 61* (CYR61) and *ankyrin repeat domain 1* (ANKRD1) by real-time PCR. Indeed, we observed induction of these genes 24 hours after APAP treatment, which reflected the expected activation of YAP (Figure 3D). To confirm YAP-dependence on the target gene induction and to exclude YAP-independent effects on gene expression, we performed additional rescue experiments. For this, the expression of YAP was silenced by RNA-interference (RNAi) in APAP-treated cells followed by the measurement of YAP target genes. According to our hypothesis, YAP inhibition partly abolished the APAP-dependent induction of CYR61 and ANKRD1 (Figure 3E).

Together, this data demonstrates that APAP predominantly acts on the Hippo pathway effector YAP in a bimodal manner. Upon APAP treatment an immediate phosphorylation/inactivation of YAP is followed by its de-phosphorylation/activation.

### APAP controls YAP phosphorylation via ROS and AKT

Based on previous data, we hypothesized a mechanistic connection between APAP-induced *reactive oxygen species* (ROS) (Barbier-Torres et al., 2017; Shuhendler et al., 2014) and AKT-driven YAP phosphorylation (Basu et al., 2003; Romano et al., 2010). This mechanistic link was investigated at early time points after APAP treatment (up to 3 hours) to exclude unspecific effects caused by APAP at later time points (e.g., due to cell toxicity).

Western immunoblotting showed that both YAP and AKT were phosphorylated early after APAP incubation (Figure 4A), which was in accordance with our previous findings (Figure 3C). In parallel, a ROS detection assay revealed that APAP induced prominent ROS activity comparable to the known ROS inducers *hydrogen peroxide* (H_2_O_2_) and *tert-butyl hydroperoxide* (TBHP) in hepatocellular cells (Figure 4B). Moreover, treatment of cells with ROS inducers demonstrated a clear (with H_2_O_2_) or moderate (with TBHP) induction of AKT and YAP phosphorylation after 1 and 2 hours, respectively (Figures 4C-D, quantification in Supplementary Figure S5A-B). This data indicated that APAP regulated ROS activity, which itself controlled YAP activity.

**Figure 4.**
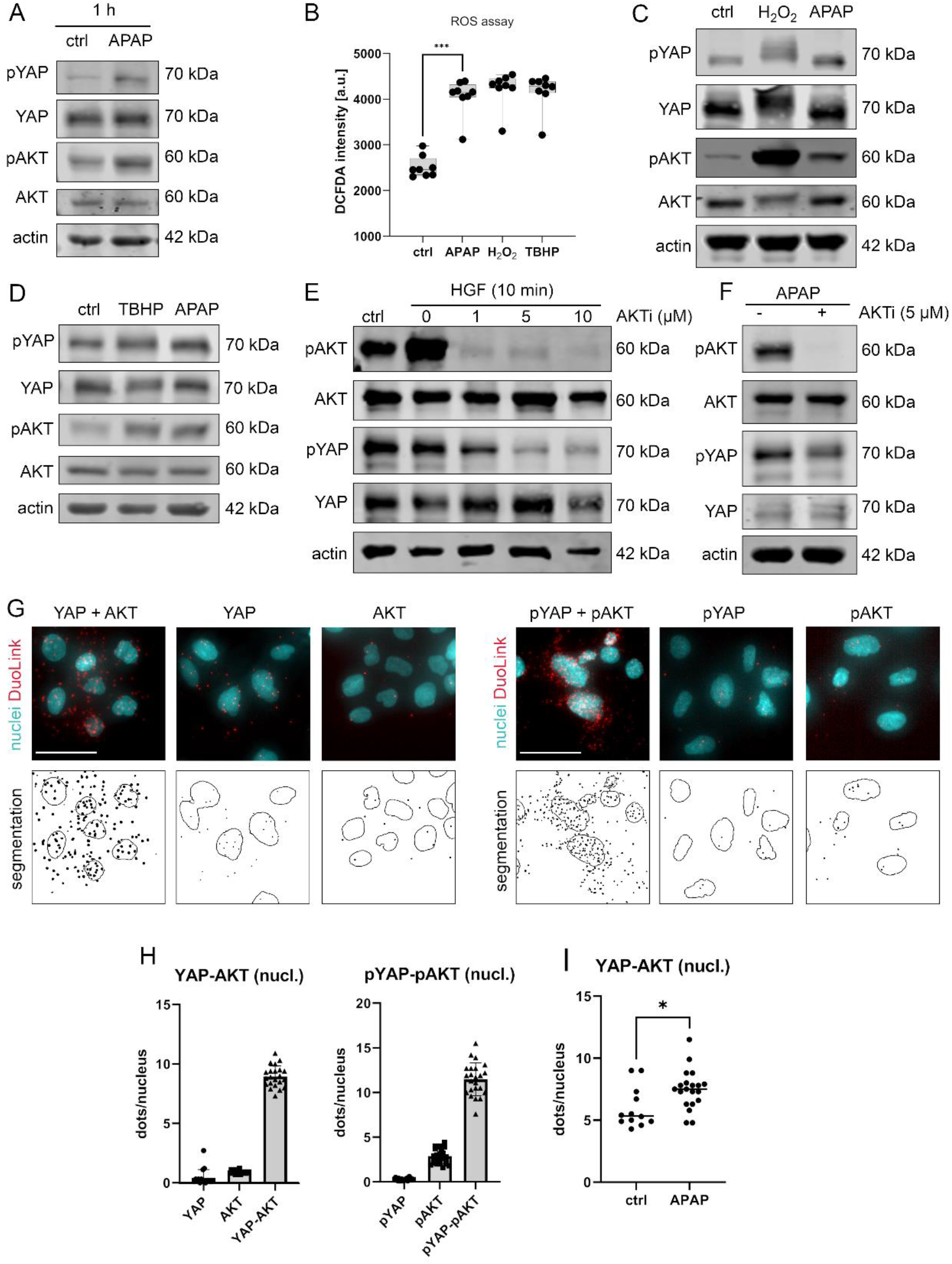
APAP controls nuclear YAP enrichment via induction of ROS and AKT. **A.** Western immunoblot analysis of pYAP, YAP, pAKT, and AKT after APAP (10 mM) administration or after treatment with PBS in Hep3B cells after one hour. The abundance of pYAP and pAKT is induced after APAP treatment in comparison to untreated cells (ctrl); n=2. **B.** Spectrophotometric ROS measurement after APAP (10 mM), hydrogen peroxide (H_2_O_2_,2 mM) and tert-butyl hydroperoxide (TBHP, 300 µM) treatment in Hep3B cells after 6 hours (n=8 technical replicates, one out of two biological replicates shown). APAP as well as H_2_O_2_ and TBHP induce ROS formation in living cells. Statistical test: one-way ANOVA with Geisser-Greenhouse correction (adjusted p-val=0.0001). Whiskers depict min and max of the dataset. **C.** Western immunoblot analysis of pYAP, YAP, pAKT, and AKT after H_2_O_2_ (2 mM) and APAP (10 mM) treatment in Hep3B cells for 1 hour. APAP, as well as H_2_O_2_ induce YAP and AKT phosphorylation compared to untreated cells (ctrl); n=4. **D.** Western immunoblot analysis of pYAP, YAP, pAKT and AKT dynamics after TBHP (300 µM) and APAP (10 mM) treatment of Hep3B cells for 2 hours. APAP and TBHP induce AKT and YAP phosphorylation as compared to respective controls (ctrl). **E.** Western immunoblot analysis of pAKT, AKT, pYAP and YAP after HGF treatment (10 ng/µl) of Hep3B cells after 10 minutes. The cells were pre-treated with an AKT inhibitor VIII (AKTi, 1 µM, 5 µM and 10 µM) and starved for 3 hours prior the HGF administration. Data illustrates that AKT inhibition prevents HGF-induced AKT phosphorylation as well as YAP phosphorylation. **F.** Western immunoblot of pAKT, AKT, pYAP and YAP after APAP (10 mM) administration for 1 hour. Cells were starved in FCS-free medium and pre-treated with AKTi (5 µM) before APAP treatment for 3 hours. Results demonstrate that AKT inhibition reduces YAP phosphorylation early after APAP treatment. **G.** PLA of the protein combinations YAP/AKT and pYAP/pAKT. Red dots indicate interactions between YAP/AKT and pYAP/pAKT, respectively. DAPI-stained nuclei are depicted in cyan. Single YAP, AKT, pYAP, and pAKT stains serve as assay controls. *Bottom row*: segmentation of the nuclei (empty circles) and dots (black spots). One out of two representative experiment shown. Scale bar: 50 µm. **H.** Quantification of the interactions between YAP/AKT and pYAP/pAKT in the nucleus (experiment shown in G). One symbol represents one image (nYAP =17, AKT=16, YAP-AKT = 22, pYAP = 24, pAKT=25, pYAP-pAKT=23). **I.** Quantification of a PLA experiment - number of YAP-AKT interaction units per nuclei of untreated (ctrl, n=12) and APAP treated cells (10 mM, for 2 hours, n=20). APAP treatment induces nuclear interaction mode between YAP and AKT. Statistical test: unpaired two-tailed parametric t-test (p-val= 0.0155). For A, C, D, E and F actin served as loading control.

To confirm the rapid and AKT-dependent phosphorylation of YAP, liver cells were treated with *hepatocyte growth factor* (HGF), which activates/phosphorylates AKT kinases *via* binding the receptor c-MET within 10 minutes (Xiao et al., 2001). HGF administration led to a clear phosphorylation of AKT but not YAP, which might be caused due to saturation effects (no further YAP phosphorylation possible under given culture conditions) (Figure 4E). However, simultaneous inhibition of AKT kinase activity by the specific inhibitor AKTi (Adlung et al., 2017; Barnett et al., 2005), not only abolished AKT phosphorylation but also reduced YAP phosphorylation (Figure 4E). As expected, upon AKTi administration concomitantly with APAP, the phosphorylation of both AKT and YAP decreased (Figures 4F, Supplementary Figure S5C).

Since our alternative PDE model predicted that the nuclear YAP phosphorylation is crucial for protein shuttling and its activity (Figure 2C-E), we hypothesized that AKT and YAP not only physically interact in the cytoplasm but also in cell nuclei. Indeed, a *proximity ligation assay* (PLA) showed that AKT and YAP, as well as the respective phosphorylated isoforms, interacted in the nucleus (Figure 4G-H). Although, interaction between AKT/YAP and pAKT/pYAP was also detectable in the cytoplasm, a prominent protein colocalization took place in the nuclear compartment (Figure 4G). Importantly, image-based quantification revealed that the interaction between nuclear YAP and AKT increased upon 6 hour treatment with APAP (Figure 4I), indicating an APAP-dependent shift of the nuclear phosphorylation and interaction rate, as predicted by the PDE model.

In summary, our data shows that APAP affects YAP activity and shuttling behavior by enhancing cellular ROS followed by nuclear AKT/YAP binding and YAP phosphorylation.

### Sequential activation of ROS, AKT, and Hippo/YAP in mouse livers after APAP intoxication

The results of our mathematical model (Figure 2) and the *in vitro* experiments (Figure 3-4) demonstrated a distinct sequence of molecular events that control YAP activity after APAP exposure: ROS induction is followed by AKT activation and YAP phosphorylation/inactivation (early events - up to 6 hours). This is followed by phase, where YAP is de-phosphorylated and transcriptionally active (late events - 24 to 48 hours).

To confirm this *in vivo*, mice were injected with a hepatotoxic dose of APAP (300 mg/kg) and liver tissues were collected up to 16 days (in total at 9 time points). Subsequently, liver specimens were subjected to expression profiling and results were investigated regarding the activity of gene expression in gene signatures specific for ROS, AKT, and Hippo/YAP activity (Han et al., 2008; Marco et al., 2017; Y. Wang et al., 2018). The experimental results showed that gene expression of ROS, AKT, and Hippo/YAP target genes in livers were altered and prominently activated between 6 hours and 2 days after APAP treatment (Figure 5A-C). To compare the temporal dynamic of the gene signature expression, z-scores of genes in the signature per time point were summarized and normalized to the number of genes in the signature. As indicated by the results from the cell culture experiments, the expression index illustrated a specific order of signature activation starting with ROS (6-12 hours), followed by AKT (12 hours to 1 day) and Hippo/YAP signature genes (1-2 days) (Figure 5D).

**Figure 5.**
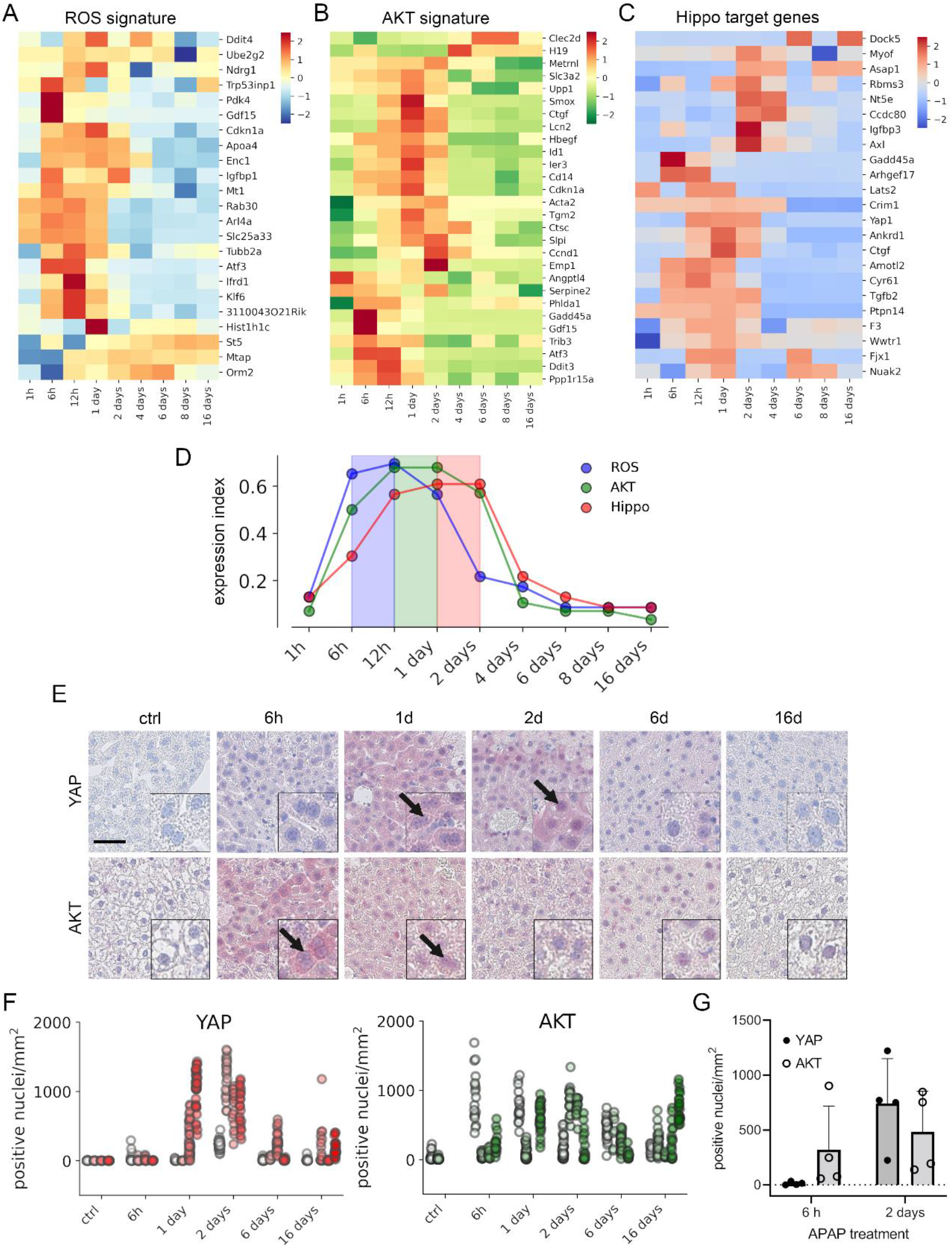
APAP stimulates the sequential activation of ROS, AKT and YAP in vivo. Gene expression profiling over time of mouse livers after APAP treatment (A-C) for up to 16 days (3-5 animals/group, 300 mg/kg). Abundance of gene signatures characteristic for the activity of ROS (A; consisting of 23 genes (Han et al., 2008)), AKT (B; consisting of 28 genes (Marco et al., 2017)) and YAP (C; consisting of 23 genes (Y. Wang et al., 2018)) were analyzed. In A-C gene expression values are z-score normalized. **D.** Summarized gene expression scores (expression index) over time illustrate the timely order of ROS (max values between 6 and 12 hours), AKT (max values between 12 hours and 1 day) and YAP (max values between 1 and 2 days) target gene signatures after APAP treatment. **E.** Mouse liver tissue sections stained for YAP (top tow) and total AKT (bottom row) under control condition and at 6 hours and 1, 2, 6, and 16 days after APAP treatment (300 mg/kg). Arrows indicate high YAP and AKT positivity in the nucleus. Scale bar: 50 µm. **F.** Automatic quantification of nuclei with YAP and AKT positivity in mouse liver tissues. Each dot represents the count of stained nuclei in one image (a tile, 1 mm^2^) for YAP (n=733) and AKT (n=520) tiles. **G.** Quantification of YAP and AKT nuclear positivity 6 hours and 2 days after APAP treatment. One dot represents average count of positive nuclei in one animal (n=4).

To confirm the findings from the gene expression analysis, we performed immunohistochemical staining of liver tissues isolated at five time points after APAP treatment. Staining for total YAP illustrated not only a general increase of YAP positivity in the cytoplasm of surviving hepatocytes, but also its prominent nuclear accumulation after 1 to 2 days (Figure 5E, arrows). Moreover, nuclear AKT enrichment was already detectable 6 hours after APAP treatment. Interestingly, nuclear AKT accumulation in hepatocytes remained high for the rest of the experiment compared to untreated mice (Figure 5E, arrows).

Because of a high mouse-to-mouse variability, we decided to objectify these results. For this, the immunohistochemical staining of YAP and AKT was quantitatively analyzed using machine learning (random forest) and image analysis methods (Figures 5F). For this, images (733 for YAP and 516 for AKT) were unbiasedly selected from all animals employed in experiment and were analyzed with an algorithm, which was trained to detect positively stained nuclei outside of the necrotic areas and the rims of the tissue (Supplementary Figure S6; for detailed information see Materials and Methods). The quantification results showed prominent nuclear YAP positivity 1 to 2 days after APAP administration, while emerging nuclear AKT was already detectable at 6 hours and maintained during the experiment (Figure 5F). The shift from exclusive nuclear AKT to nuclear YAP was illustrated by direct comparison of the time points 6 hours and 2 days after APAP injection (Figure 5G).

To sum up, we confirm the sequential order of APAP-induced events that cause YAP activation in mouse liver tissue.

## Discussion

In this study we aimed to investigate the different properties of the Hippo pathway effectors YAP and TAZ under *in vitro and in vivo* conditions that resembled the situation after drug-induced liver damage. For this, we developed a cell-based model system that allowed the systematic and comparative tracing of spatial YAP/TAZ dynamics upon stimulation with APAP, which is one of the most widely used analgesic with high potential of liver toxicity. By integrating time-resolved experimental data, computational modeling, image analysis tools as well as confirmatory *in vivo* results, we could draw several important conclusions on the role of YAP and TAZ under physiological conditions and in liver cells upon DILI.

Several mathematical modeling approaches have been previously applied to explain the dynamic behavior of the Hippo pathway under physiological or pathological conditions in different organisms. For example, computational modeling was used to explain how Hippo signaling cross-talks with other signaling pathways *via* protein interactions (Labibi et al., 2020; Romano et al., 2014). Other studies exclusively focused on biophysical/biochemical regulation of the Hippo pathway to investigate how YAP/TAZ activity relates to mechanical input information (Ege et al., 2018; Eroumé et al., 2021; Gou et al., 2018; Scott et al., 2021; Sun et al., 2016).

We decided to use PDE modeling, which allowed us to discover spatial aspects of signaling dynamics and to use live-cell image data for parametrizing the Hippo signaling steady state models. By doing so, we compared PDE model simulations of different model topologies with the experimentally acquired data for YAP as well as TAZ and selected a model topology which best corresponded to data generated *in vitro*. The selected alternative Hippo pathway model is based on experimental observations and parametrized within the considered boundaries of experimental evidence (see supplementary data). The obtained model parameters are results of the best model fit obtained, and allow reproducing the described model behavior. However, since the model fitting is based on steady state data only, and concentration observations are in arbitrary units, the parameter values are subject to structural non-identifiability. Nevertheless, this does not affect the qualitative conclusions drawn from the models for the investigated biological process.

The results of our PDE modeling approach strongly suggested that the nuclear phosphorylation reaction of YAP/TAZ was necessary and sufficient for the reproduction of their subcellular distribution patterns *in vitro* (i.e., the halo-like nuclear distribution of YAP and TAZ). This observation is of importance since the conventional view on the Hippo pathway topology represents sequential phosphorylation steps in the cytoplasm. Our PDE modeling-based finding would therefore extend the current understanding of Hippo pathway regulation since it adds the cell nucleus as a pivotal spatial compartment in the regulation of Hippo/YAP/TAZ signaling.

This conclusion is supported by our experimental findings illustrating that LATS1/2 are highly expressed in cell nuclei and that the interaction between phosphorylated LATS1/2 and YAP as well as between AKT and YAP are predominantly detectable in nuclei. Importantly, previously published experimental and computational approaches support our findings. For example, nuclear LATS1/2 controls YAP phosphorylation in the context of NF2/Merlin-deficiency (Li et al., 2014). In addition, the combination of quantitative photo-bleaching experiments and modeling illustrated that nuclear export is a crucial reaction for the subcellular YAP distribution (Ege et al., 2018). Combining these insights on nuclear export mechanisms with our findings on nuclear YAP phosphorylation strongly suggests that both processes closely cooperate in the efficient regulation of the Hippo pathway effectors. Lastly, our model does not explicitly exclude phosphorylation reaction in the cytoplasm; however, it suggests the necessity to re-evaluate the canonical Hippo pathway scheme. The appeal for the re-evaluation and extension of the canonical Hippo pathway, e.g., by combining models on nuclear phosphorylation and nuclear transport, would further broaden our understanding on the dynamic spatial behavior of complex signaling pathways (Ege et al., 2018; Shreberk-Shaked & Oren, 2019).

To our knowledge, a direct mathematical comparison of YAP and TAZ regarding their spatial distribution has not been performed, yet. Using the PDE model, we comparatively investigated YAP and TAZ distribution and identified kinetic parameters which discriminate between both species. Specifically, the observed localization differences (nuclear exclusion *vs.* nuclear localization) between YAP and TAZ might be caused by distinct phosphorylation rates of YAP and TAZ. Nuclear/cytoplasmic shuttling dynamics for YAP and/or TAZ have been investigated previously (Ege et al., 2018; Plouffe et al., 2018). However, our model predicts that the phosphorylation efficiency for TAZ must be higher than for YAP to achieve the observed differential localization of YAP and TAZ.

By utilizing our image analysis approach for the detection of minor changes in the subcellular distribution of YAP and TAZ, we showed a dynamic shuttling response of YAP (and to a much lesser extent of TAZ) upon APAP treatment. Although, APAP-dependent activation of YAP was described in the literature (Poudel et al., 2021), the underlying molecular mechanisms were poorly understood. Our results demonstrated that APAP-induced YAP shuttling is not the result of an unspecific cellular response but is due to a sequential order of molecular events that include ROS induction (Ghallab et al., 2022; Shuhendler et al., 2014), followed by AKT (in)activation (Koundouros & Poulogiannis, 2018) and nuclear YAP (de)phosphorylation. Time-dependent APAP-induced ROS activation was recently supported by a study by Ghallab et al., which showed transient induction of oxidative stress in mice during 8 hours after APAP overdose with a peak at 2 hours (Ghallab et al., 2022). Interestingly, we also observed a biphasic YAP regulation: YAP phosphorylation/inactivation at early time points (up to 3 hours) and its dephosphorylated/active at later time points (>24 hours). Especially, the functionally important activation of YAP, which is associated with the induction of cell proliferation, is well reflected by the temporal activation of YAP in murine tissues upon APAP administration after 24 to 48 hours.

Importantly, several cellular mechanisms might simultaneously contribute to the APAP-dependent activation of YAP. For example, the direct impact of APAP on MST1/2 kinases, which are central Hippo pathway constituents, are one additional process that may contribute to the observed effects on YAP (Fan et al., 2016). Moreover, the serine-/threonine-kinase AKT could control the Hippo/YAP axis at different levels. For example, a physical interaction of AKT and YAP in the nucleus (shown here) or the cytoplasm might directly control YAP phosphorylation (Basu et al., 2003). Alternatively, AKT can phosphorylate MST2 and therefore indirectly contribute to YAP activity (Romano et al., 2014). Thus, APAP probably controls YAP in a multi-modal manner to achieve an optimal cellular response and it is therefore likely that this multimodal process is part of cell-protective mechanism that counteracts the massive loss of hepatocytes upon APAP overdose (Holland et al., 2021; Shuhendler et al., 2014).

## Materials and Methods

### In Vitro Experiments

#### Cell culture and the establishment of genetically modified cells

The hepatocyte-derived cell lines Hep3B (#ACC93, DSMZ, Braunschweig, Germany), HLF, HepG2, and SNU182 (ATCC/LGC Standards, Wesel, Germany) were cultured in MEM, DMEM and RPMI medium, respectively. The media were supplemented with 10% FCS and 1% penicillin-streptomycin (Sigma-Aldrich, Taufkirchen, Germany). Cells were maintained at 37°C in a 5% CO_2_ atmosphere. Hep3B cells were transduced with vectors carrying cDNAs for human histone H2B, YAP, and TAZ genes, fused with mCerulean, mVenus, and mCherry genes, respectively. The plasmid pRRLN-EF1α-mVenus-YAP was kindly provided by Prof. Dr. Alexander Loewer (TU Darmstadt), pLentiPGK-mCerulean-H2B was purchased (Addgene, Watertown, USA; No. 90234). The pRRLN-EF1α-d2mCherry-TAZ was cloned. Correctness of vectors was verified by end-to-end sequencing.

Vectors were stably integrated into the Hep3B cells using lentiviral particles. For this, HEK293 cells were transfected with lentiviral vectors, packaging (psPAX2, Addgene, Watertown, USA; No. 12260), and envelope plasmids (pMD2.G, Addgene, Watertown, USA; No. 12259) using polyethylenimine and incubated for 40 hours. Subsequently, virus particles in the cell culture medium were collected and used for infections. The transduced cells were selected for stable vector integration using geneticin (0.6 µg/ml, pRRLN-EF1α-mVenus-YAP), hygromycin (0.6 µg/ml, pLentiPGK-mCerulean-H2B), and blasticidin (2 µg/ml, pRRLN-EF1α-mVenus-YAP). Cells were frequently checked for mycoplasma contamination and authenticated by *short tandem repeat* (STR) analysis (DSMZ, Braunschweig, Germany).

The APAP stock solution (100 mM) was freshly prepared by dissolving 0.3 g APAP crystalline powder in 20 ml phosphate-buffered saline (PBS, GE Healthcare, Solingen, Germany) under heating (42°C) and stirring (Sigma-Aldrich, St. Louis, USA). The APAP solution was filtered (Millex-GS filters, 0.22 µm; Merck Millipore Ltd., Carrigtwohill, Ireland) and immediately administered to cells to a final concentration of 10 mM. *Tert-butyl hydroperoxid* (TBHP, Luperax, Sigma-Aldrich) with 70 wt. % in H_2_O was applied to cells to a final concentration of 300 µM for 2 hours prior protein extraction. *Hydrogen peroxide* (H_2_O_2_) 30% (Rotipuran, Carl Roth GmbH, Karlsruhe, Germany), was applied to a final concentration of 2 mM for 1 hour.

AKT1/2 phosphorylation was inhibited with InSolution™ Akt Inhibitor VIII (1-10 µM, Merck KGaA, Darmstadt, Germany) in FCS-free medium for 3 hours prior to treatment with HGF (10 ng/µl).

#### Gene silencing by RNA interference (RNAi)

RNAi was performed to inhibit YAP gene expression. For this, 2×10^5^ cells were seeded per well in a 6 cm-well plate and incubated overnight. Oligofectamine (Invitrogen, Carlsbad, CA, USA) and Opti-MEM (Gibco Life Technologies Ltd., Paisley, UK) were used for transfection of the siRNA (final concentration: 5 nM) according to the manufacturer’s instructions. A random sequence oligonucleotide was used as a control treatment (control siRNA: ctrl siRNA). The following siRNA sequences were used: YAP siRNA #1 (5’-3’): CCA CCA AGC UAG AUA AAG A-dT-dT and YAP siRNA #2 (5’-3’): GG UCA GAG AUA CUU CUU AA-dT-dT, and ctrl siRNA (5’-3’): UGG UUU ACA UGU CGA CUA A. YAP siRNA #1 and #2 were pooled to achieve an optimal gene knockdown. Twenty-four hours after siRNA transfection, cells were treated with APAP (10 mM) for 24 hours.

#### Live cell imaging

Stably transduced Hep3B cells were seeded on 24-well glass-bottom black wall plates (MoBiTec GmbH, Goettingen, Germany) in a phenol red-free RPMI 1640 medium (Gibco/Life Technologies Corporation, Paisley, UK). Dynamic protein localization in living cells was measured using confocal laser scanning microscope (Nikon A1R on an inverted Nikon Ti2 microscope), which was supplemented with an on-stage incubator (TokaiHit) to maintain 37°C and 5% CO_2_ at all stages of the experiments. Selected images were acquired with additional Nikon AxR microscope system.

Three lasers were employed: 445 nm (mCerulean, PMT detector), 514 nm (mVenus, GaAsP detector), and 561 nm (mCherry, GaAsP detector) to obtain highly resolved live-cell images. Two channels – 445 nm and 514 nm - were acquired simultaneously. The employed pinhole size was 17.9 µm (445 nm: 1.5 a.u., 514 nm: 1.3 a.u., 561 nm: 1.2 a.u.). Images were acquired using a galvano-scanner of size 512 x 512 px (field of view 0.64 x 0.64 mm, pixel resolution of 1.24 µm). The objectives Plan Apo λ 20x (NA 0.75, working distance 1 mm, FOV 0.64 x 0.64 mm) or Plan Apo λ 40x (NA 0.95, working distance 0.21 mm, FOV 0.435 x 0.435 mm) were used.

#### Western immunoblotting

Western immunoblotting was performed as previously described (Weiler et al., 2017). In brief, for total protein fraction preparation, cultured cells were harvested using 1x Cell Lysis Buffer (Cell Signaling Technology, Danvers, USA) supplemented with 1x Protease Inhibitor Mix G (Serva, Heidelberg, Germany) and 1x PhosSTOP (Roche Diagnostics Deutschland GmbH, Mannheim, Germany). For protein fractionation, the NE-PER Nuclear and Cytoplasmic Extraction kit was used (Thermo Scientific, Rockford, USA). The following antibodies were used for Western immunoblotting experiments: YAP XP (1:1’000, Cell Signaling Technology, Danvers, USA, #14074, RRID:AB_2650491), pYAP (1:400, Cell Signaling Technology, #4911, RRID:AB_2218913), pAKT (1:1’000, Cell Signaling Technology, #4060, RRID:AB_2315049), AKT (1:1’000, Cell Signaling Technology, #9272, RRID:AB_329827), β-actin (1:10’000, MP Biomedicals, Solon, USA, #08691001, RRID:AB_2335127), LATS1 (1:200, Santa Cruz Biotechnology, Dallas, USA, sc-398560), LATS2 (1:200, Santa Cruz Biotechnology, sc-515579), and pLATS1/2 (1:500, Cell Signaling Technology, #8654, RRID:AB_10971635) and Cytochrome P450 2E1 (1:500, Novus Biologicals, Centennial, USA, # NVP1-85367, RRID:AB_11021447). PARP (1:10’000, Cell Signaling Technology, #9542, RRID:AB_2160739), β-tubulin (1:200, Santa Cruz Biotechnology, sc-5274, RRID:AB_2288090) and GAPDH (1:10’000, Merck Millipore, Darmstadt, Germany, ab2302, RRID:AB_10615768) were used as loading controls.

Western Blot detection and quantification was performed using the Odyssey-CLx Infrared Imaging system with the ImageStudio software (LI-COR Biosciences, Bad Homburg, Germany). Phosphorylated protein band intensity was measured and normalized to total protein concentrations. Equal amount of protein was loaded for each line, as measured by the Bradford reagent (Millipore Sigma, Saint Louis, USA). Raw unedited image files and uncropped blots are available as source data files of this manuscript.

#### RNA isolation, reverse transcription, and quantitative real-time PCR (qPCR)

RNA was isolated using Extract me kit (Blirt, Gdańsk, Poland) according to the manufacturer’s protocol. Reverse transcription was performed using the RevertAid kit (Thermo Scientific). Semi-quantitative real-time PCR reactions were set up using the primaQuant 2x qPCR-SYBR-Green-Mastermix (Steinbrenner Laborsysteme, Wiesenbach, Germany) and analyzed with the QuantStudio 3 real-time PCR system (Applied Biosystems, Thermo Fischer Scientific, Singapore). The following cycling conditions were applied: 95°C for 15 min, followed by 40 cycles of 95°C for 15 s, and 60°C for 60 s. Product specificity was confirmed by melting curve analysis (95°C for 15 s, 60°C for 30 s, 60–95°C with 0.5°C/s).

The mRNA levels were normalized to *glyceraldehyde-3-phosphate dehydrogenase* (GAPDH), *60S ribosomal protein L41* (RPL41), *serine/arginine-rich splicing factor* (SRSF4), and *β2-microglobulin* (B2M). The following primers for human cDNAs were used: YAP-for: 5’-CCT GCG TAG CCA GTT ACC AA-3’; YAP-rev: 5’-CCA TCT CAT CCA CAC TGT TC-3’; ANKRD1-for: 5’-AGT AGA GGA ACT GGT CAC TGG-3’; ANKRD1-rev: 5’-TGG GCT AGA AGT GTC TTC AGA T-3’; CYR61-for: 5’-AGC CTC GCA TCC TAT ACA ACC-3’; CYR61-rev: 5’-TTC TTT CAC AAG GCG GCA CTC-3’; GADPH-for: 5’-CTG GTA AAG TGG ATA TTG TTG CCA T-3’; GAPDH-rev: 5’-TGG AAT CAT ATT GGA ACA TGT AAA CC-3’; RPL41-for: 5’-AAA CCT CTG CGC CAT GAG AG-3’; RPL41-rev: 5’-AGC GTC TGG CAT TCC ATG TT-3’; SRSF4-for: 5’-TGC AGC TGG CAA GAC CTA AA-3’; SRSF4-rev: 5’- TTT TTG CGT CCC TTG TGA GC-3’; B2M-for: 5’- CAC GTC ATC CAG CAG AGA AT-3’; B2M-rev: 5’- TGC TGC TTA CAT GTC TCG AT-3’.

#### In situ proximity ligation assay (PLA)

The DuoLink *in situ* PLA was performed according to the manufacturer’s instructions (Sigma-Aldrich). Briefly, cells were seeded on glass coverslips and prior to APAP administration cells were grown under FCS-free conditions for one day, then incubated with APAP (10 mM) or PBS. After treatment, cells were washed three times with 2 mM MgCl_2_ in PBS and fixed with 4% paraformaldehyde for 10 min at room temperature. Fixed cells were washed four times with PBS for 5 min, permeabilized with 0.2% Triton X-100 in PBS for 5 min at room temperature, and washed again twice with PBS for 5 min. Subsequently, cells were blocked with Blocking solution (Sigma-Aldrich) for 30 min at room temperature and incubated with the following primary antibodies diluted in Antibody Diluent (Sigma-Aldrich) overnight at 4°C: anti-YAP (1:25, Santa Cruz Biotechnology, sc-271134, RRID:AB_10612397), anti-AKT (1:200, Cell Signaling Technology, #9272, RRID:AB_329827), anti-pYAP (1:200, Cell Signaling Technology, #4911, RRID:AB_2218913), anti-pAKT (1:200, Cell Signaling Technology, #4051, AB_331158), and anti-pLATS1/2 (1:200, Cell Signaling Technology, #8654, RRID:AB_10971635). Subsequently, cells were washed twice with Wash Buffer A (Sigma-Aldrich) and incubated with pre-diluted rabbit PLUS and mouse MINUS probes (Sigma-Aldrich) in Antibody Diluent for 1 hour at 37°C. After incubation, cells were washed twice with Wash Buffer A and then incubated with ligation solution (Sigma-Aldrich) for 30 min at 37°C. After ligation, samples were again washed twice with Wash Buffer A for 2 min at room temperature and incubated with amplification solution, and detection reagent Orange (Sigma-Aldrich) for 100 min at 37°C. Finally, cells were washed twice with Wash Buffer B (Sigma-Aldrich) for 10 min at room temperature, once with 0.01X Wash Buffer B for 1 min at room temperature and coverslips were mounted on the slide with DAPI Fluoromount-G mounting medium (SouthernBiotech, Birmingham, USA).

Fluorescence images were captured using an inverted Nikon Ti2 microscope with Nikon S Plan Fluor ELWD 40x NA 0.60 objective in a widefield fluorescence mode using Lumencor Sola SE II lamp. Images were captured in DAPI and TRITC channels (460 nm and 580 nm) with Nikon DS-Qi2 monochrome camera (image size 2404×2404 px, pixel resolution of 0.18 µm/px).

#### Reactive oxygen species (ROS) activity measurement

Measurements of the intracellular ROS levels were performed using the DCFDA/H2DCFDA cellular ROS assay kit (Abcam, Amsterdam, Netherlands) according to the manufacturer’s protocol. In brief, cells were seeded on white clear-bottom 96-well plate (Corning, Corning, USA) and incubated overnight. Cells were treated with 10 mM APAP, 2 mM H_2_O_2_, or 300 µM TBHP (H_2_O_2_ and TBHP served as positive controls for ROS induction) for 6 hours in FCS-free cell culture medium. Fluorescence was measured using a microplate reader (FluoStar Omega, BMG Labtech GmbH, Ortenberg, Germany). Buffer solution without cells served as a background control.

### In Vivo Experiments and Sample Analyses

#### Housing and treatment of mice and induction of acute liver injury by acetaminophen

Male C57BL6/N mice (8 to 10-week old) were bought from Janvier Labs (Janvier Labs, Le Genest-Saint-Isle, France). Animals were housed under 12 hours light/dark cycles at controlled ambient temperature of 25°C with free access to water and were fed *ad libitum* with a standard diet (Ssniff, Soest, Germany) before starting the experiments. Induction of acute liver injury with APAP was done as previously described (Schneider et al., 2021). Briefly, the mice were fasted overnight, then challenged with a dose of 300 mg/kg APAP intraperitonealy. APAP was dissolved in warm PBS with an application volume of 30 ml/kg. Control group was treated with PBS only. The mice were fed *ad libitum* after APAP administration. All animals were included for further analyses. All experiments were approved by the local animal welfare committee (LANUV, North Rhine-Westphalia, Germany, application number: 84-02.04.2016.A279).

#### Liver tissue sample collection, processing, and staining

Tissues were collected time-dependently after APAP injection from the left liver lobe. The tissues were fixed and embedded in paraffin as previously described (Ghallab et al., 2016). YAP and AKT immunostaining were performed using 4 µm-thick paraffin-embedded tissue sections. For immunohistochemistry, an anti-YAP antibody (1:50, Cell Signaling Technology, #14074, RRID:AB_2650491) and anti-AKT (1:50, Cell Signaling Technology, #9272, RRID:AB_329827) were used. Embedded tissue sections were pre-treated with a heat induced epitope retrieval (HIER) method (pH 6, DAKO, Hamburg, Germany). As secondary antibody anti-rabbit Polymer-AP (Enzo Life Sciences, Farmingdale, USA, ENZ-ACC110-0150) was used. Detection was performed with Permanent AP (Zytomed Systems GmbH, Berlin, Germany). Following staining, the whole slides were digitally documented using a slide scanner (Aperio AT2, Leica Mikrosysteme Vertrieb GmbH, Wetzlar, Germany).

#### Expression profiling and bioinformatics

RNA isolation and gene array analysis were performed as published before (Campos et al., 2020) and bioinformatic analysis was done as described (Holland et al., 2021). The gene expression data is available under ArrayExpress accession number GSE167032. The expression data was applied to three known signatures that are informative for ROS activity (Han et al., 2008), AKT activity (Marco et al., 2017) and YAP/TAZ activity (Y. Wang et al., 2018). Due to the high number of AKT signature genes, only genes whose response to APAP treatment was larger than fold change of 2 (compared to control animals) were considered. The expression data was z-score normalized and clustered with seborn cluster map python module (v0.11.0). The signature score was obtained by summarizing z-scored expression values at the given time point if the z-scored value was greater than 0.5. The summarized expression values were normalized to the number of genes in the signature.

### Computational Methods

#### Analysis of the live-cell images

The confocal images of the living cells were analyzed in a high-throughput manner using ImageJ (v1.53f51) platform (Rueden et al., 2017). Images were first manually selected with respect to the quality criteria: images with insufficient sharpness or with artefacts were discarded from the analysis. The image processing pipeline was based on Weka segmentation (v3.3.1) (Arganda-Carreras et al., 2017) of foreground and background areas, and subsequent thresholding, object detection, and counting. After object detection and counting, the respective masks were overlaid on the initial images to acquire YAP and TAZ intensity values for nuclei and cytoplasm. Mean pixel intensity from nuclear areas of the cells was divided by the mean pixel intensity of cytoplasmic regions, thus obtaining a *nuclear-to-cytoplasmic ratio* (NCR).

#### Analysis of PLA images

PLA slides were analyzed using ImageJ (v1.53f51). First, nuclei and dots were classified using the Weka segmentation algorithm (v3.3.1) (Arganda-Carreras et al., 2017), thresholded, and counted. The pseudo-cytoplasmic (ring-shaped) area was created using the ImageJ’s binary mask option *dilate* for 30 iterations on nuclear masks to obtain nuclear and cytoplasmic area for a comparative analysis. The thresholded nuclei or cytoplasmic masks were overlaid with detected dots to obtain the information on the subcellular localization of the protein interaction.

#### Analysis of IHC images

Stained tissue samples were digitalized using Aperio slide scanner with 40x magnification and pixel resolution of 0,253 µm/px (Aperio AT2, Leica Mikrosysteme Vertrieb GmbH, Wetzlar, Germany). The selected time-points (6 hours and 1, 2, 6, 16 days) and control treatment were quantified using a pipeline, which consisted of python scripts (modules PIL v5.3.0, matplotlib v2.2.3), ASAP software (v1.6, https://github.com/computationalpathologygroup/ASAP) and Ilastik software (v1.3.3) (Berg et al., 2019).

First, digital images of the tissue sections were divided in tiles (1 mm^2^, 400×400 pixels), which were binned (to reduce processing time and storage load) using ASAP and python modules. Some images were excluded due to staining and/or scanning artifacts. Tiles, which displayed tissue for at least 50% of their area, were kept for further processing. Machine learning model, which was based on a random forest algorithm, was trained on a selected set of training tiles for YAP or AKT using Ilastik software. The algorithm was trained to detect positively stained nuclei, excluding necrotic areas and staining artefacts. Ilastik software was further used to export probability maps, which were possessed with ImageJ. In ImageJ, probability maps were thresholded and positive nuclei were counted and normalized to the area of the tile, which was occupied by the tissue.

#### Partial differential equation (PDE) modeling

Spatial modeling aimed at describing the variations in space of fluorescently labelled YAP and TAZ distribution within living Hep3B cells. The mathematical model is based on a system of *partial differential equations* (PDEs). PDE modeling and parameter estimation was performed with the Spatial Model Editor (SME) software (v1.2.1) and sme-contrib (v0.0.14) python module (https://spatial-model-editor.github.io/). SME is graphical user interface-based model editing and simulation software compatible with systems biology markup language (SBML) standards. Models were simulated using simple Forward Time Centered Space (FTCS) solver.

In the demonstrated models all reactions were defined by first-order kinetics (for PDEs see supplementary material). For the parameter estimation, the model behavior was evaluated at steady state, i.e. the model was simulated until the concentration distribution did not change over time. Parameter estimation was performed using the particle swarm algorithm (20 particles, 200 iterations) to minimize a cost function consisting of the weighted sum of two terms: the squared per pixel differences between model and the target image, and the sum of squares of species concentration rates of change. In total 200 fitted parametrizations for each tested model were generated. The parameter space for the optimization algorithm was defined based on published data (see supplementary material). The mathematical models were uploaded to the BioModels repository under the model identifier number MODEL2202080001.

#### Statistics

Statistical analysis was performed using GraphPad Prism 9.2.0. Statistical tests are indicated in figure legends. Error bars depict standard deviation. Significance levels are as follows: *p-value ≤ 0.05, **p-value ≤ 0.01, ***p-value ≤ 0.001.

## Acknowledgements

We thank the Nikon Imaging Center at Heidelberg University (BioQuant), especially Dr. Ulrike Engel and Dr. Christian Ackermann, for supporting confocal imaging experiments. We also acknowledge the LSDF2 (URZ) for data storage. Further we thank Prof. Dr. Alexander Loewer for providing the YAP-mVenus vector.

## Author Contributions

**Lilija Wehling**: study concept and design, live cell image data acquisition and analysis, and interpretation of data, PDE modeling study design, modeling and parameter estimation and manuscript writing. **Kai Breuhahn**: study concept, design and supervision, interpretation of data and manuscript writing. **Sven Sahle** : spatial model editor development, PDE modeling design, supervision, manuscript writing. **Liam Keegan**: spatial model editor development, revision of the manuscript. **Paula Fernández-Palanca**: performed Western blotting and PLA experiments and analysis, revision of the manuscript. **Jennifer Schmitt**: performed cell culture and Western blotting experiments, and provided technical support. **Reham Hassan**: performed mouse experiment and tissue collection, revision of the manuscript. **Ahmed Ghallab** and **Jan G. Hengstler**: mouse study concept, design and data acquisition, revision of the manuscript. **Peter Schirmacher**: pathological evaluation of tissues and revision of the manuscript. **Ursula Kummer**: PDE modeling study concept and revision of the manuscript.

## Data availability

The mathematical models generated and analyzed during the current study are available in the *BioModels* repository (https://www.ebi.ac.uk/biomodels/) under accession number MODEL2202080001. The PDE simulation software is available over *Github* (https://spatial-model-editor.github.io/). The gene expression data analyzed during the current study is available in the *Gene Expression Omnibus (GEO)* repository under accession number GSE167032 (https://www.ncbi.nlm.nih.gov/geo/).

## Ethics approval

This study was performed according to animal welfare committee of The Ministry for Environment, Agriculture, Conservation and Consumer Protection of the State of North Rhine-Westphalia (LANUV, North Rhine-Westphalia, Germany, application number: 84-02.04.2016.A279).

## Supplementary Information for

**This PDF file includes:**

Supplementary text (PDE model equations)

Figures S1 to S6

Tables S1 to S6

SI References

### PDE model equations

In the computational model the equations which are simulated in cytoplasm or nucleus follow two-dimensional reaction diffusion equation. Initial conditions for the PDE are arbitrary since only steady state solutions are considered. The equations in TAZ model are equivalent to YAP model equations.

Here, described in general terms as follows:

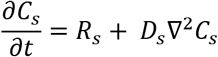

where

- *C_s_* is the concentration of species *S* at position (*x*, *y*) and time *t*
- *R_s_* is the reaction term for species *S*
- *D_s_* is the diffusion constant for species *S*
- ∇^2^ *C_s_* is the Laplacian of species *S*, which can be rewritten as

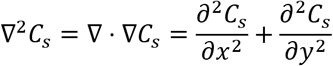

The partial differential equations for the dynamics of YAPn/pYAPn (nuclear fraction) and YAP/pYAP (cytosolic fraction) proteins in the alternative model in the nucleus or cytoplasm are presented as follows:

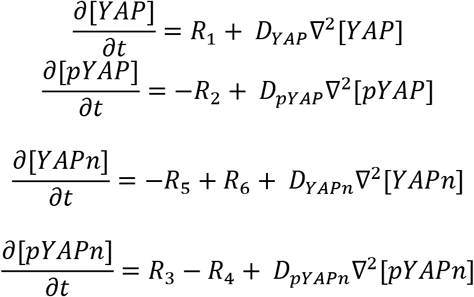

Where reaction rates *R* are defined as follows:

R_1_ as translation:

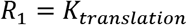
R_2_ as degradation:

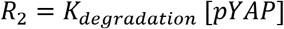
R_5_ as phosphorylation:

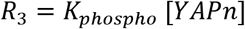
R_6_ as dephosphorylation:

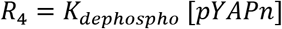

The transport reactions for YAP and pYAP are defined as flux densities across the nuclear membrane that depend on the respective concentrations; these are translated into Neumann type interface conditions for the inner boundary of the cytoplasm and the outer boundary of the nucleus by the simulator software.

- flux density of YAP import

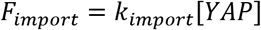
- flux density of YAP export

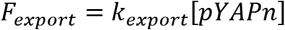

Boundary conditions on the outer membrane of the cytoplasm are “zero-flux” Neumann type for all variables.

### Model Parameters

**Table S1.** Canonical model parameters.

| Parameter | Symbol | YAP model | TAZ model | Unit |
| --- | --- | --- | --- | --- |
| Diffusion | $D_{YAP}$<br>$D_{YAPn}$ | 0.0391776 | 0.00812799 | $\mu\text{m}^2/\text{s}$ |
| Diffusion phosphorylated | $D_{pYAPn}$<br>$D_{pYAP}$ | 0.433785 | 0.501116 | $\mu\text{m}^2/\text{s}$ |
| Phosphorylation | $K_{phospho}$ | 3.2462731384467 | 9.2660885020866 | $\text{s}^{-1}$ |
| Dephosphorylation | $K_{dephospho}$ | 2.4380091829261 | 2.9721728333365 | $\text{s}^{-1}$ |
| Translation | $K_{translation}$ | 0.013295743091699 | 0.02329254213393 | a.u./s |
| Degradation | $K_{degradation}$ | 0.0081058910608829 | 0.0090941179957213 | $\text{s}^{-1}$ |
| Nuclear import | $K_{import}$ | 9.7229134833679e-16 | 8.1106772583928e-16 | $\mu\text{m}/\text{s}$ |
| Nuclear export | $K_{export}$ | 9.5119620861314e-17 | 6.1050891990604e-17 | $\mu\text{m}/\text{s}$ |

**Table S2.** Alternative model parameters.

| Parameter | Symbol | YAP model | TAZ model | Unit |
| --- | --- | --- | --- | --- |
| Diffusion | $D_{YAP}$<br>$D_{YAPn}$ | 9.84707 | 9.14393 | $\mu\text{m}^2/\text{s}$ |
| Diffusion phosphorylated | $D_{pYAPn}$<br>$D_{pYAP}$ | 1.04518 | 1.00732 | $\mu\text{m}^2/\text{s}$ |
| Phosphorylation | $K_{phospho}$ | 0.028839815689838 | 0.10286291836829 | $\text{s}^{-1}$ |
| Dephosphorylation | $K_{dephospho}$ | 0.09121073484801 | 0.10517790271802 | $\text{s}^{-1}$ |
| Translation | $K_{translation}$ | 0.009403085138566 | 0.0030932046019853 | a.u./s |
| Degradation | $K_{degradation}$ | 0.0094299166245222 | 0.0097940321843269 | $\text{s}^{-1}$ |
| Nuclear import | $K_{import}$ | 6.5845623444598e-15 | 9.5850453479782e-15 | $\mu\text{m}/\text{s}$ |
| Nuclear export | $K_{export}$ | 7.8819897236132e-15 | 4.5876638224922e-15 | $\mu\text{m}/\text{s}$ |

**Table S3.** Parameters of the YAP-to-TAZ transition model.

| Parameter | Symbol | Value | Unit |
| --- | --- | --- | --- |
| Diffusion | $D_{YAP}$<br>$D_{YAPn}$ | 10 | $\mu\text{m}^2/\text{s}$ |
| Diffusion phosphorylated | $D_{pYAPn}$<br>$D_{pYAP}$ | 1 | $\mu\text{m}^2/\text{s}$ |
| Phosphorylation | $K_{phospho}$ | - | $\text{s}^{-1}$ |
| Dephosphorylation | $K_{dephospho}$ | - | $\text{s}^{-1}$ |
| Translation | $K_{translation}$ | 0.01 | a.u./s |
| Degradation | $K_{degradation}$ | 0.004 | $\text{s}^{-1}$ |
| Nuclear import | $K_{import}$ | 7.5e-15 | $\mu\text{m}/\text{s}$ |
| Nuclear export | $K_{export}$ | 1.6e-15 | $\mu\text{m}/\text{s}$ |

**Table S4.** Parameter range for particle swarm algorithm of the canonical YAP/TAZ model.

| Parameter | Symbol | Range (min, max) | Unit |
| --- | --- | --- | --- |
| Diffusion | $D_{YAP}$ | (0.001, 5) | $\mu\text{m}^2/\text{s}$ |
| | $D_{YAPn}$ | | |
| Diffusion phosphorylated | $D_{pYAPn}$<br>$D_{pYAP}$ | (0.001, 1) | $\mu\text{m}^2/\text{s}$ |
| Phosphorylation | $K_{phospho}$ | (0.1, 10) | $\text{s}^{-1}$ |
| Dephosphorylation | $K_{dephospho}$ | (0.1, 10) | $\text{s}^{-1}$ |
| Translation | $K_{translation}$ | (1e-3, 0.1) | a.u./s |
| Degradation | $K_{degradation}$ | (1e-3, 1e-2) | $\text{s}^{-1}$ |
| Nuclear import | $K_{import}$ | (1e-16, 1e-15) | $\mu\text{m}/\text{s}$ |
| Nuclear export | $K_{export}$ | (1e-17, 1e-15) | $\mu\text{m}/\text{s}$ |

**Table S5.** Parameter range for particle swarm algorithm of the alternative YAP/TAZ model.

| Parameter | Symbol | Range (min, max) | Unit |
| --- | --- | --- | --- |
| Diffusion | $D_{YAP}$<br>$D_{YAPn}$ | (1, 10) | $\mu\text{m}^2/\text{s}$ |
| Diffusion phosphorylated | $D_{pYAPn}$<br>$D_{pYAP}$ | (1, 1.5) | $\mu\text{m}^2/\text{s}$ |
| Phosphorylation | $K_{phospho}$ | (0.01, 0.5) | $\text{s}^{-1}$ |
| Dephosphorylation | $K_{dephospho}$ | (0.01, 0.5) | $\text{s}^{-1}$ |
| Translation | $K_{translation}$ | (1e-3, 0.01) | a.u./s |
| Degradation | $K_{degradation}$ | (1e-3, 0.01) | $\text{s}^{-1}$ |
| Nuclear import | $K_{import}$ | (1e-16, 1e-14) | $\mu\text{m}/\text{s}$ |
| Nuclear export | $K_{export}$ | (1e-16, 1e-14) | $\mu\text{m}/\text{s}$ |

**Table S6.** Parameter values from the literature.

| Parameter | Value | Source |
| --- | --- | --- |
| Hepatocyte cell volume | 10-11 L | (1) |
| Hepatocyte nucleus volume | $5 \cdot 10^{-13}$ | (1) |
| Nuclear export | 10-100 s | (2) |
| Nuclear import | 50 s | (2) |
| Diffusion rate YAP/TAZ | $19 \mu\text{m}^2/\text{s}$ | (2) |

## SI Figures

**Fig. S1.**
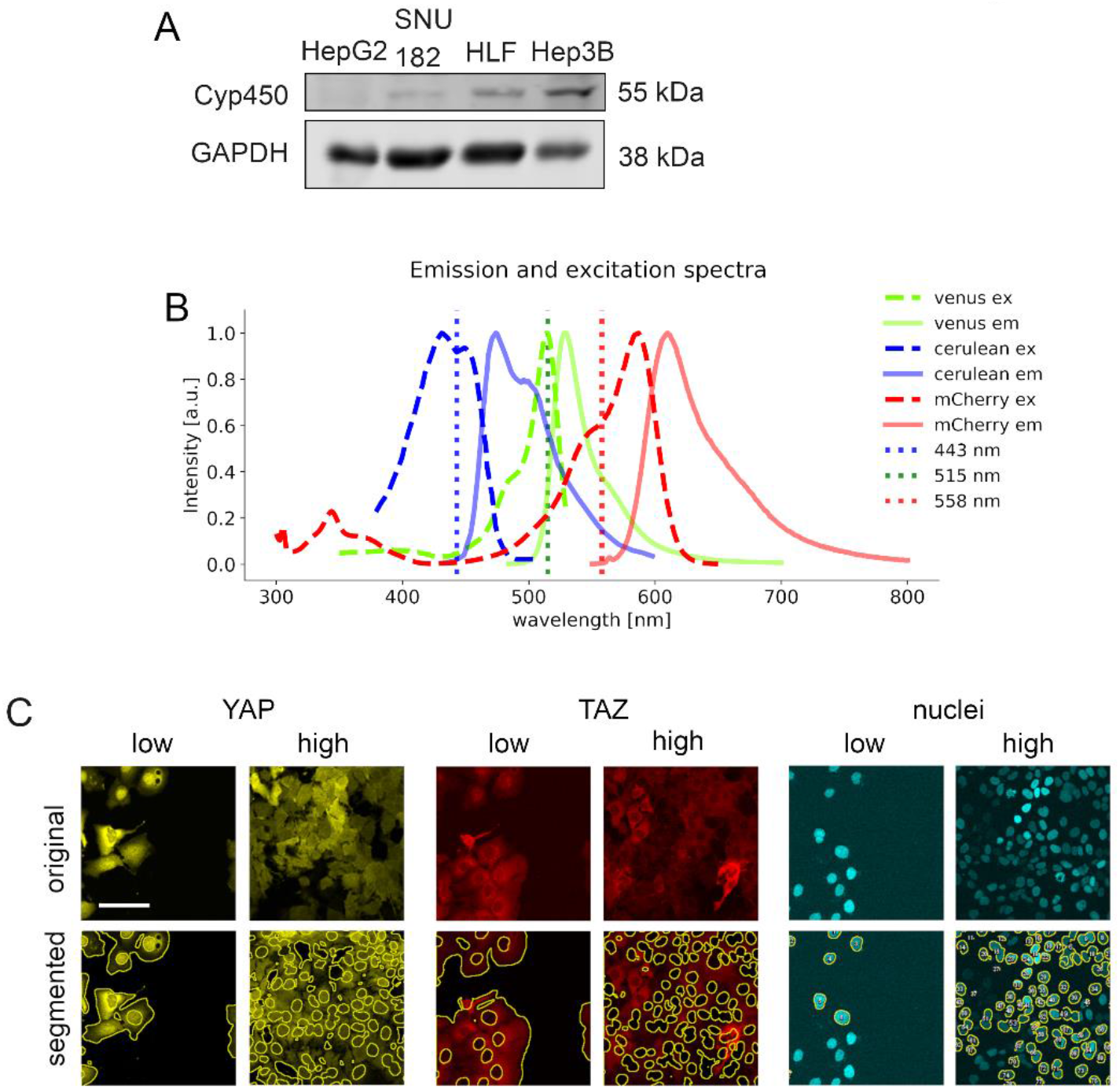
*In vitro* model establishment. **A.** Western immunoblotting of cell lines HepG2, SNU182, HLF and Hep3B, which were tested for the abundance of cytochrome P450 2E1 (Cyp450), a liver enzyme which metabolizes APAP. Hep3B cells show a prominent Cyp450 expression. GAPDH serves as a loading control. **B.** Emission and excitation spectra of Venus, Cerulean and mCherry fluorophores. Dotted line represents the wavelength of lasers, which were used for the confocal microscopy setup. **C.** Exemplary segmentation results of the image analysis pipeline for nuclei, YAP and TAZ localization under low and high cell density conditions. Yellow lines represent outlines of the segmented areas for the cytoplasmic positivity (either YAP-Venus or TAZ-mCherry) or nuclear marker (H2B-Cerulean). Images are gamma corrected (γ=0.5). Scale bar: 100 µm.

**Fig. S2.**
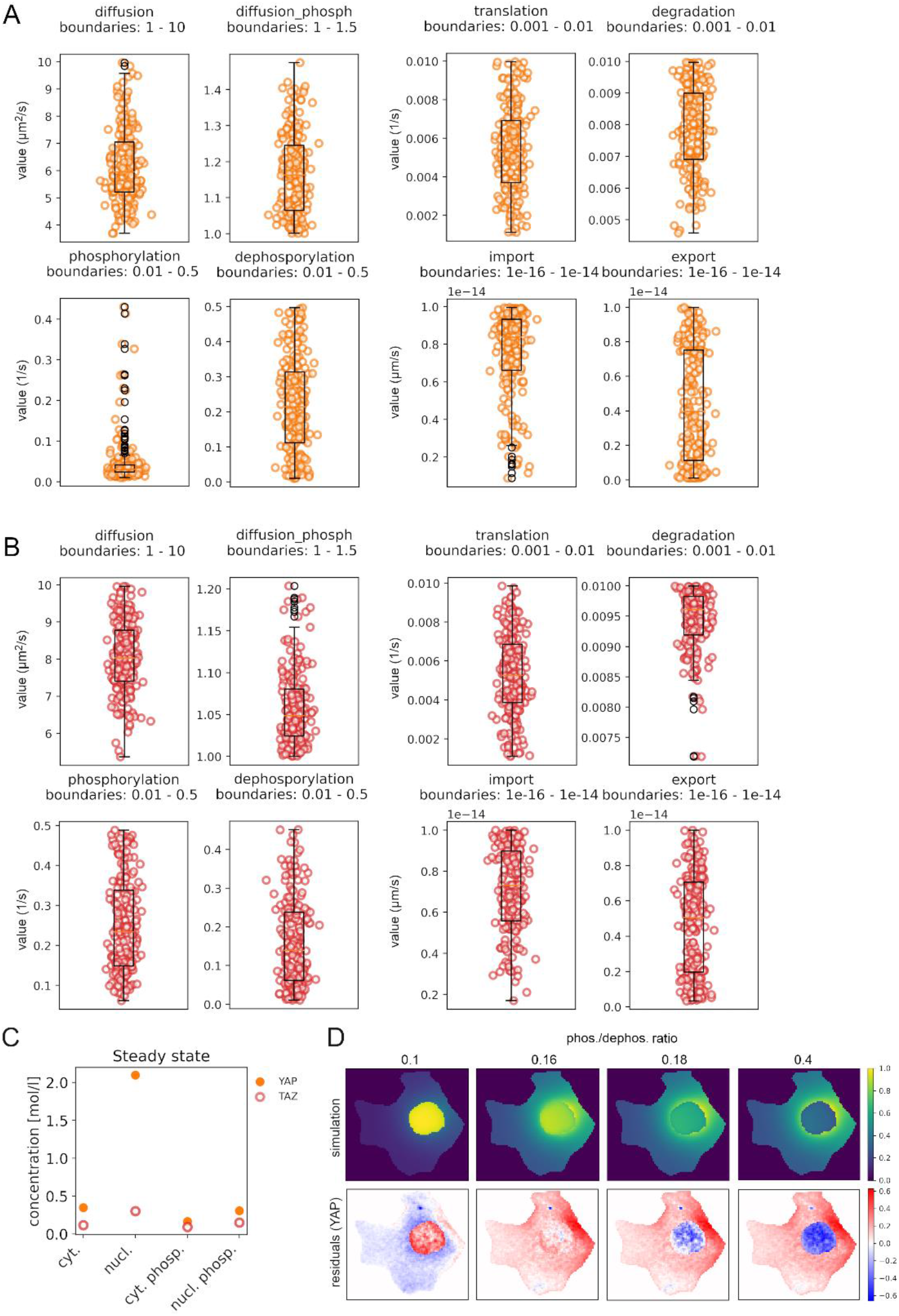
Parametrizing the computational models. Distribution of parameter values for each fitted parameter of alternative Hippo pathway models for YAP (A) and TAZ (B) proteins. Here, 200 models were obtained by parameter estimation method with particle swarm algorithm (200 iterations, 20 particles). Obtained parameters are plotted within the min and max values of the evaluated parameter space boundaries. **C.** Steady state values of YAP/TAZ and pYAP/pTAZ concentrations in nuclei and cytoplasm of an exemplary alternative model for YAP and TAZ (model simulation depicted in Figure 2D). **D.** Alternative model simulations with respect to changes in phosphorylation to dephosphorylation ratio (phos./dephos. rato). Increasing phos./dephos. ratio drives YAP protein from mainly nuclear localization to cytoplasmic, mirroring the differences between YAP and TAZ localization under low cell density conditions in a steady state.

**Fig. S3.**
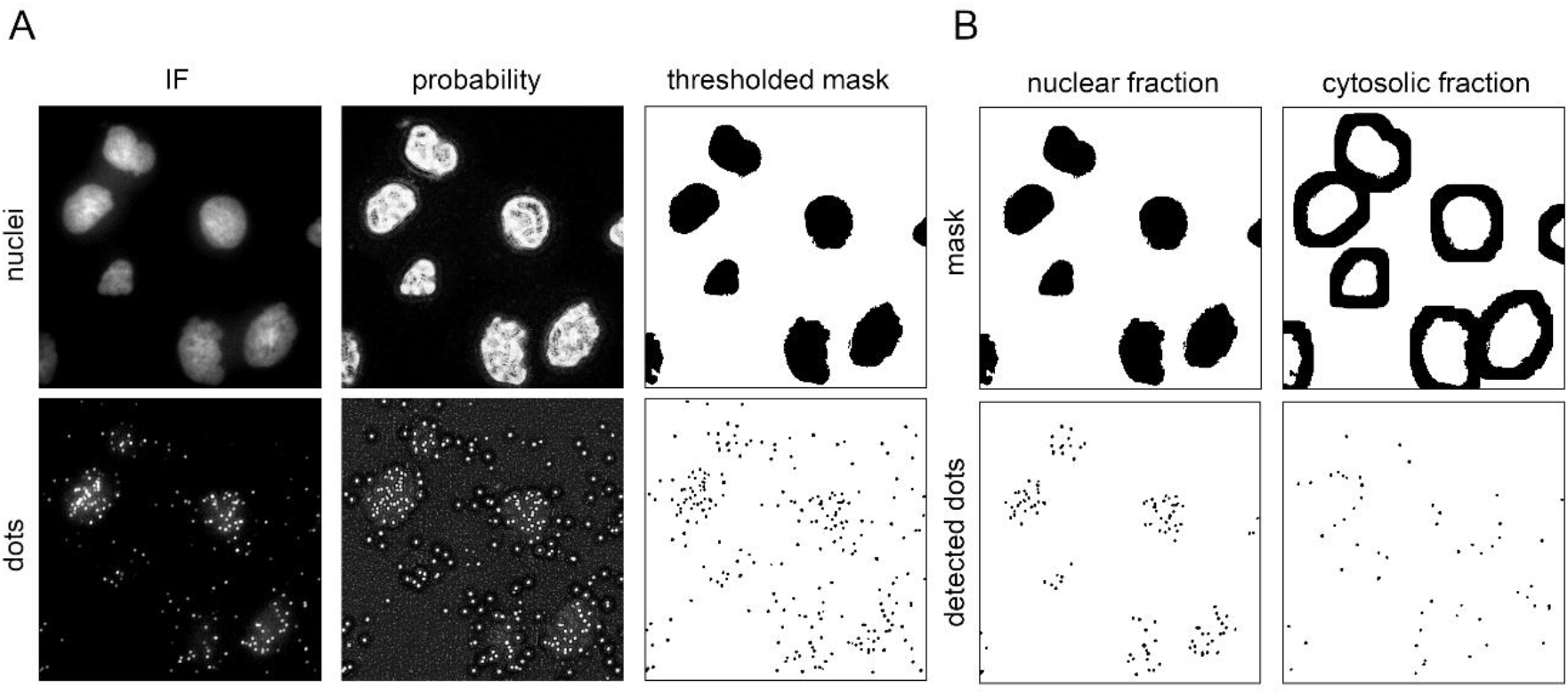
Data analysis pipeline for PLA quantification. **A.** First steps of the image analysis pipeline involve generation of the probability maps by applying trained Weka segmentation model onto images of nuclei and the PLA dots. Further, by setting a threshold to the probability maps, masked images of nuclei and dots are generated. IF: immunofluorescence. **B.** The number of dots within the area of nuclear mask and cytosolic mask are counted. Cytosolic mask was obtained by expanding the nuclear area for 30 iterations using ImageJ’s dilate function. This generated a cytoplasmic area comparable to the respective nuclear area for each cell.

**Fig. S4.**
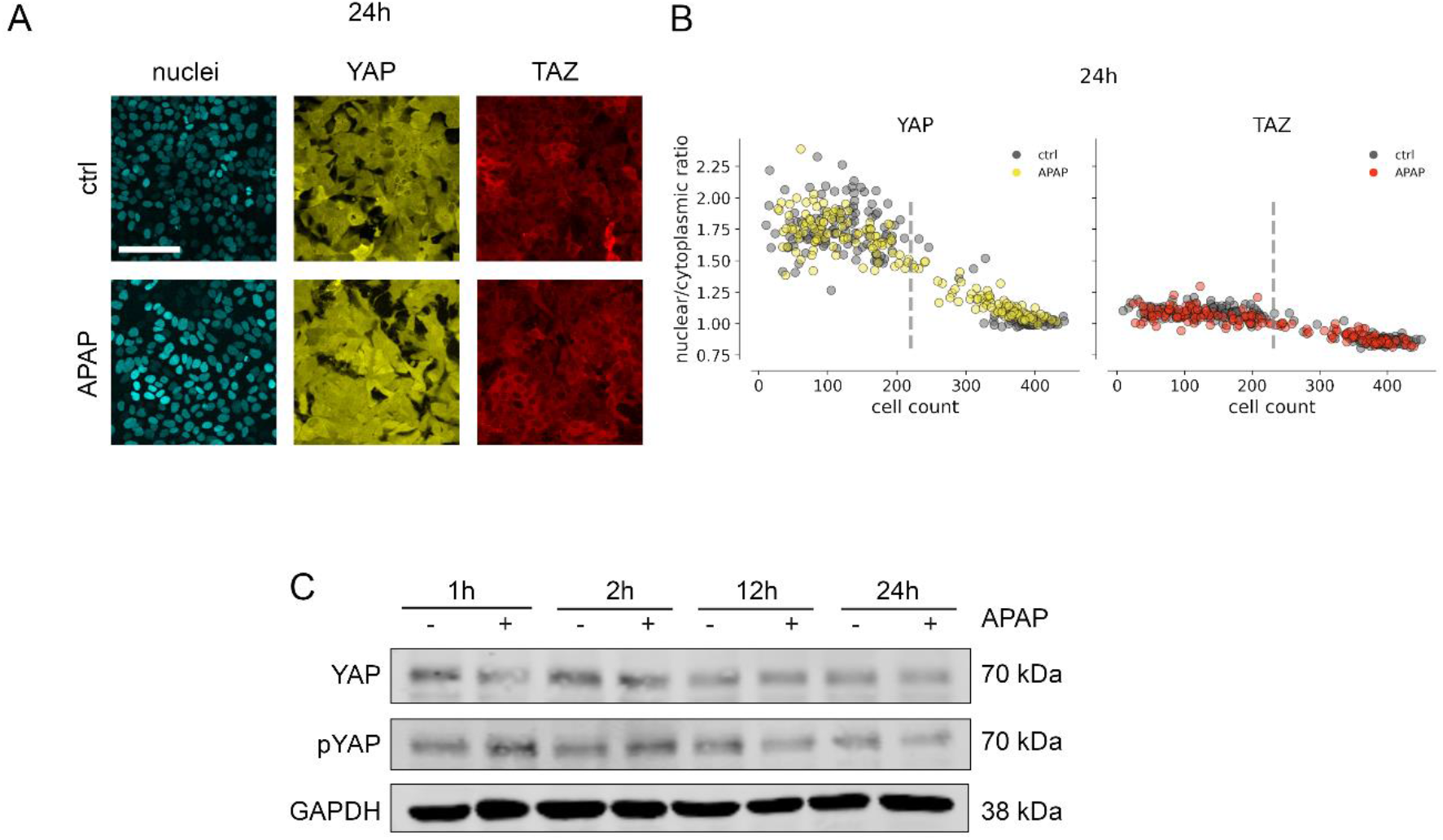
APAP treatment of hepatocellular cells induces YAP and TAZ dynamics **A.** The subcellular localization of YAP and TAZ 24 hours after and APAP 10 (mM) treatment in comparison to untreated cells (ctrl). Scale bar: 100 µm. **B.** Quantification of the YAP and TAZ NCR dynamics in relation to cell density changes 24 hours after APAP (10 mM) treatment. **C.** Time-dependent dynamics of YAP and pYAP expression after APAP (10 mM) treatment in HepG2 cells.

**Fig. S5.**
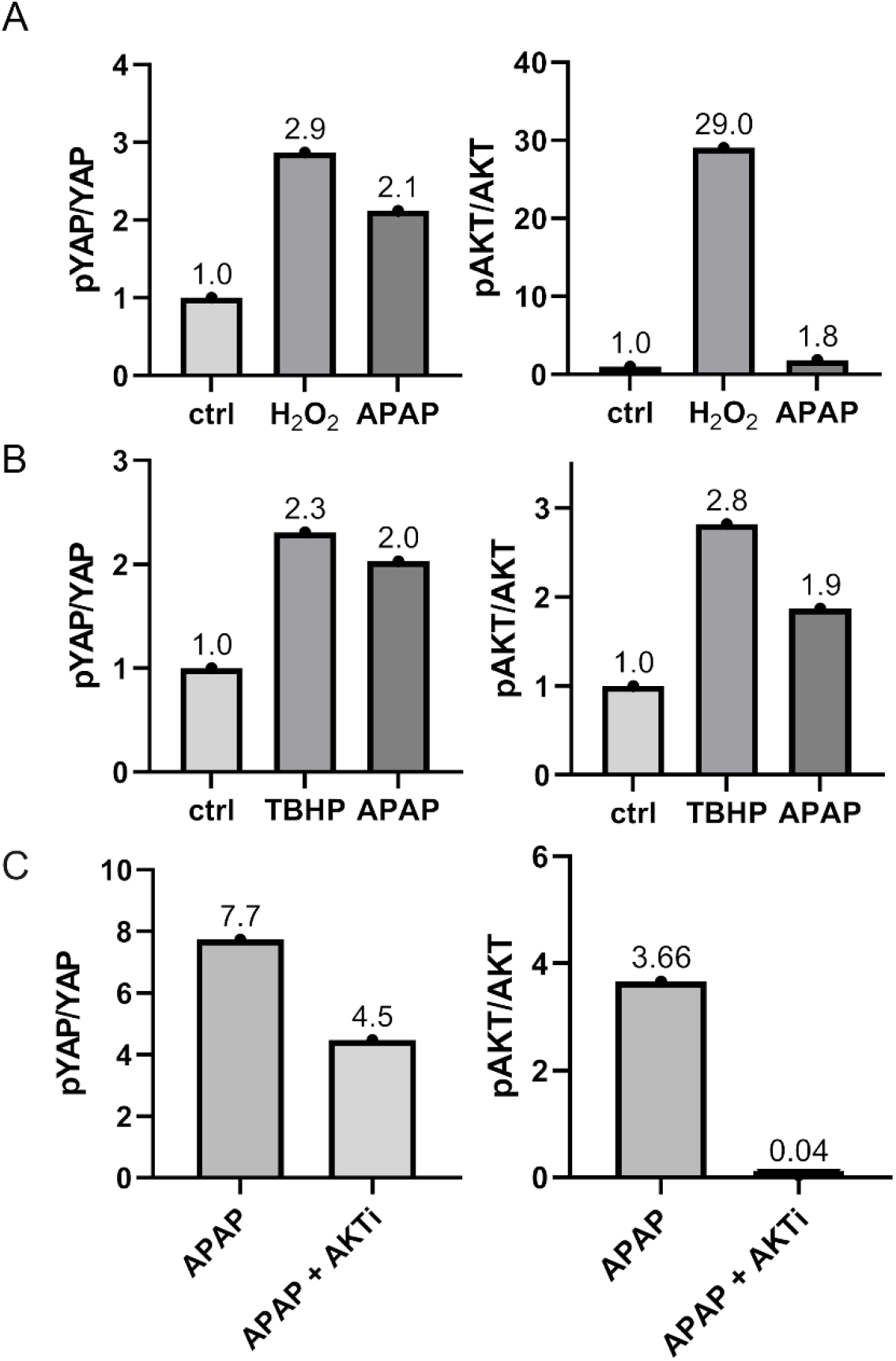
Quantification of the Western immunoblot analysis with respect to pYAP to YAP and pAKT to AKT ratio. Panel A corresponds to Figure 4C; B to Figure 4D; C corresponds to Figure 4F.

**Fig. S6.**
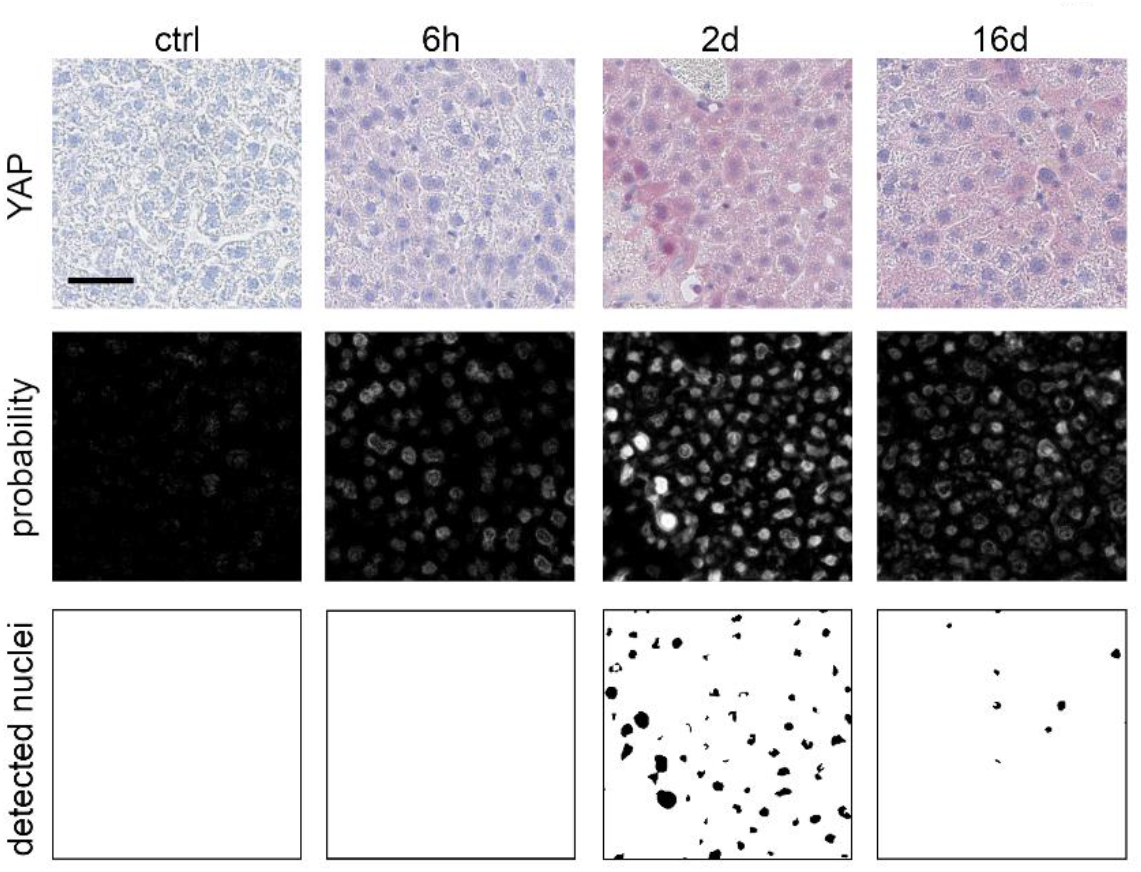
Quantification of YAP- and AKT-positive nuclei on immunohistochemically stained tissue slides. Exemplary nuclei quantification workflow for YAP-positive nuclei in mouse liver tissues. Top row: IHC stain for YAP in untreated tissue samples and after APAP treatment (300 mg/kg). The respective time after APAP treatment is indicated above. Middle row: probability maps obtained by the trained model. Bottom row: detected nuclei after applying threshold on the probability maps.

